# Human pluripotent stem cell-derived kidney organoids reveal tubular epithelial pathobiology of heterozygous *HNF1B*-associated dysplastic kidney malformations

**DOI:** 10.1101/2023.03.14.532598

**Authors:** Ioannis Bantounas, Kirsty M. Rooney, Filipa M. Lopes, Faris Tengku, Steven Woods, Leo A. H. Zeef, Shweta Y. Kuba, Nicola Bates, Sandra Hummelgaard, Katherine A Hillman, Silvia Cereghini, Adrian S. Woolf, Susan J. Kimber

## Abstract

*Hepatocyte nuclear factor 1B* (*HNF1B*) encodes a transcription factor expressed in developing human kidney epithelia. Heterozygous *HNF1B* mutations are the commonest monogenic cause of dysplastic kidney malformations (DKMs). To understand their pathobiology, we generated heterozygous *HNF1B* mutant kidney organoids from CRISPR-Cas9 gene-edited human ESCs and iPSCs reprogrammed from a family with *HNF1B*-asscociated DKMs. Mutant organoids contained enlarged malformed tubules and displayed deregulated cell turnover. Numerous genes implicated in Mendelian kidney tubulopathies were downregulated, and mutant tubules resisted the cAMP-mediated dilatation seen in controls. Bioinformatic analyses indicated abnormal WNT, calcium, and glutamatergic pathways, the latter hitherto unstudied in developing kidneys. Glutamate ionotropic receptor kainate type subunit 3 was upregulated in mutant organoids and was detected in their tubules and in fetal human DKM dysplastic epithelia. These results reveal morphological, molecular, and physiological roles for HNF1B in human kidney tubule morphogenesis and functional differentiation. They additionally suggest druggable targets to ameliorate disease.

## INTRODUCTION

The human metanephric kidney initiates at 5 weeks of gestation (Woolf and Jenkins, 2015; Woolf 2019) when metanephric mesenchyme (MM) is penetrated by the ureteric bud (UB). MM/UB crosstalk (McMahon, 2016) results in MM undergoing mesenchymal to epithelial transition to generate primitive nephrons, each differentiating into a glomerulus, a proximal tubule (PT) and a distal tubule (DT), including the loop of Henle. Waves of nephrogenesis forming glomeruli and nephron tubules occur between antenatal weeks 7 and 34, and the UB arborises into collecting ducts (CDs), each fusing with a DT. Stromal fibroblast-like cells (Wilson et al., 2021) and blood and lymphatic endothelia (Jafree et al., 2019) are found between developing kidney tubules. Capillaries invade glomerular podocyte tufts and deliver blood for filtration, and tubules modify the ultrafiltrate to make definitive urine. Dysplastic kidney malformations (DKMs) are major causes of kidney failure in children (Kohl et al., 2022) and young adults (Pepper et al., 2022). Ultrasonography detects echo-bright organs that, lacking cortical-medullary distinction (Kohl et al., 2022), manifest aberrant nephron morphogenesis and patterning accompanied by deregulated cell turnover (Winyard et al., 1996a and 1996b).

Some DKMs have genetic causes (Deng et al., 2022; Kohl et al., 2022), the commonest being heterozygous variants of *hepatocyte nuclear factor 1B* (*HNF1B*) on chromosome 17q12 (Adalat et al., 2010; Adalat et al., 2019). *HNF1B* has nine exons and encodes a homeodomain transcription factor that binds a canonical DNA sequence (Adalat et al., 2010) after dimerizing with itself or *HNF1A* (Bach et al., 1991; Lu et al., 2007; Clissold et al., 2015). *HNF1B*-associated kidney disease can vary in severity, without clear genotype-phenotype correlations (Lim et al., 2020). Some fetuses with minimal excretory function undergo termination (Humaitre et al., 2006; Duval et al., 2016). Others survive gestation but postnatally exhibit urinary electrolyte wasting, suggesting that HNF1B is required for tubular functional differentiation (Adalat et al., 2010; Adalat et al., 2019). *HNF1B* is expressed in developing human nephron epithelia and CDs (Kolatsi-Joannou et al., 2001; Haumaitre et al., 2006). Expression is maintained in mature kidney epithelia excepting podocytes, while MM and stromal cells do not express *HNF1B*. *HNF1B*-associated DKMs contain abnormal multi-layered tubules and dysmorphic glomeruli (Bingham et al., 2002; Humaitre et al., 2006; Duval et al., 2016; Nakayama et al., 2022).

Functional studies have manipulated HNF1B in kidney somatic cell lines (Chan et al., 2020; Piedrafita et al., 2021), *Xenopus* (Grand et al., 2023), and mice. Cre-mediated deletion of both *Hnf1b* alleles in mouse metanephroi deregulates expression of genes including those implicated in WNT signalling and polycystic kidney disease (PKD) (Lokmane et al., 2010; Desgrange et al., 2017; Gresh et al., 2004; Fiorentino et al., 2020). Postnatal biallelic deletion perturbs mouse kidney mitochondrial respiration (Casemayou et al., 2017), enhances fibrosis (Chan et al., 2018) and impairs regeneration (Verdeguer et al., 2010). Germline homozygous mutant mice are, however, early embryonic lethal and humans with biallelic germline mutations have not been described. Thus, the relevance of biallelic models to understand human heterozygous *HNF1B*-associated DKMs is uncertain. Recently, however, aberrant tubules were described in kidneys of mice carrying a heterozygous intron 2 splice donor site germline *Hnf1b* mutation (Niborski et al., 2021). Nevertheless, kidney gene expression patterns are not identical in humans and mice (Lindstrom et al., 2018), so applying mouse models to understand human developmental disease can still be questioned. New human experimental models are needed to overcome these limitations.

Generating kidney organoids from human pluripotent stem cells (hPSCs) is being used to understand normal and abnormal kidney development (Taguchi et al., 2014; Takasato et al., 2015; Bantounas et al 2018,2020; Woolf 2019; Rooney et al., 2021). We hypothesized that hPSC-derived kidney organoids would model human *HNF1B-*associated DKMs and would illuminate their underlying pathobiology. We therefore undertook morphological, functional, and molecular analyses of heterozygous *HNF1B* mutant organoids derived from CRISPR-Cas9 gene edited wild-type human embryonic stem cells (hESCs) or from human induced PSCs (hiPSCs) reprogrammed from peripheral blood mononuclear cells (PBMNs) of siblings with *HNF1B*-associated DKMs. Mutant organoids contained malformed nephrons and displayed deregulated proliferation and apoptosis. Numerous genes implicated in Mendelian tubulopathies were downregulated in mutant organoids which resisted cAMP-mediated tubule dilatation, present in wild-type controls. Bioinformatic analyses predicted abnormal pathways including WNT and glutamatergic signalling, hitherto unstudied in kidney development. Glutamate ionotropic receptor kainate type subunit 3 (GRIK3) was markedly upregulated in mutant organoids and was also detected in human *HNF1B*-associated DKM tubules. Our results therefore illuminate important roles for HNF1B in human kidney development and identify potentially druggable targets to treat DKMs.

## RESULTS

### Generating heterozygous *HNF1B* mutant hESCs

CRISPR/Cas9 gene editing was used to mutate *HNF1B* in MAN13 hESCs (Ye et al., 2017) (Fig. 1A). Of 21 clones isolated and sequenced, 3 were heterozygous mutants and one (IBM13-19) was studied further. This clone carried a frameshift in exon 1 (Fig. 1B) such that mutant mRNA would contain a premature stop codon 51 nucleotides downstream. Thus mutant protein would lack DNA binding and transactivation domains (Clissold et al., 2015). A wild-type clone (IBM13-08) constituted an isogenic non-mutant control line (Fig. 1B). To determine whether the mutant line had nephrogenic potential we used a 2D protocol that generates primitive kidney epithelia from control hESCs (Bantounas et al, 2018; Bantounas et al., 2021). Immunocytochemistry was undertaken for HNF1B with an antibody raised against a 111 amino acid epitope that would be disrupted by the mutation, and for epithelial cell-cell junction protein cadherin-1 (CDH1, or E-cadherin). Control cells formed CDH1+/HNF1B+ islands (Fig. 1C). The mutant line also formed CDH1+ islands but HNF1B immunoreactivity appeared attenuated (Fig. 1C). We next undertook a differentiation protocol (Fig. 1D) that induces hPSCs to form nephron-rich organoids (Bantounas et al., 2018 and 2021; Takasato et al., 2015;). Control and mutant hESCs formed organoids (Fig. 1E) of similar area (Fig. 1F) but phase contrast microscopy suggested that internal structures were larger in mutant organoids (Fig. 1G). Using exon 2 primers that would detect both non mutant and mutant mRNA, qPCR showed that *HNF1B* increased similarly in the two genotypes over the 25 day protocol (Fig. 2A). Western blotting detected HNF1B as a 63 kDa doublet in control non mutant and mutant organoids (Fig. 2B) and densitometry confirmed lower levels in those of mutants (Fig. 2C). To determine which cells expressed *HNF1B*, a BaseScope *in-situ* hybridization (ISH) probe was used that would detect both non mutant and mutant transcripts. Most control tubules expressed *HNF1B* (Fig. 2D) and in mutant organoids *HNF1B* was detected in bulky, morphologically aberrant, tubules with a lower signal in glomeruli (Fig. 2D). Using immunohistochemistry, HNF1B was detected in nuclei in tubules of control organoids (Fig. 2E) but the signal appeared attenuated and less distinctly nuclear in mutant dysplastic tubules (Fig. 2E).

**Figure 1.**
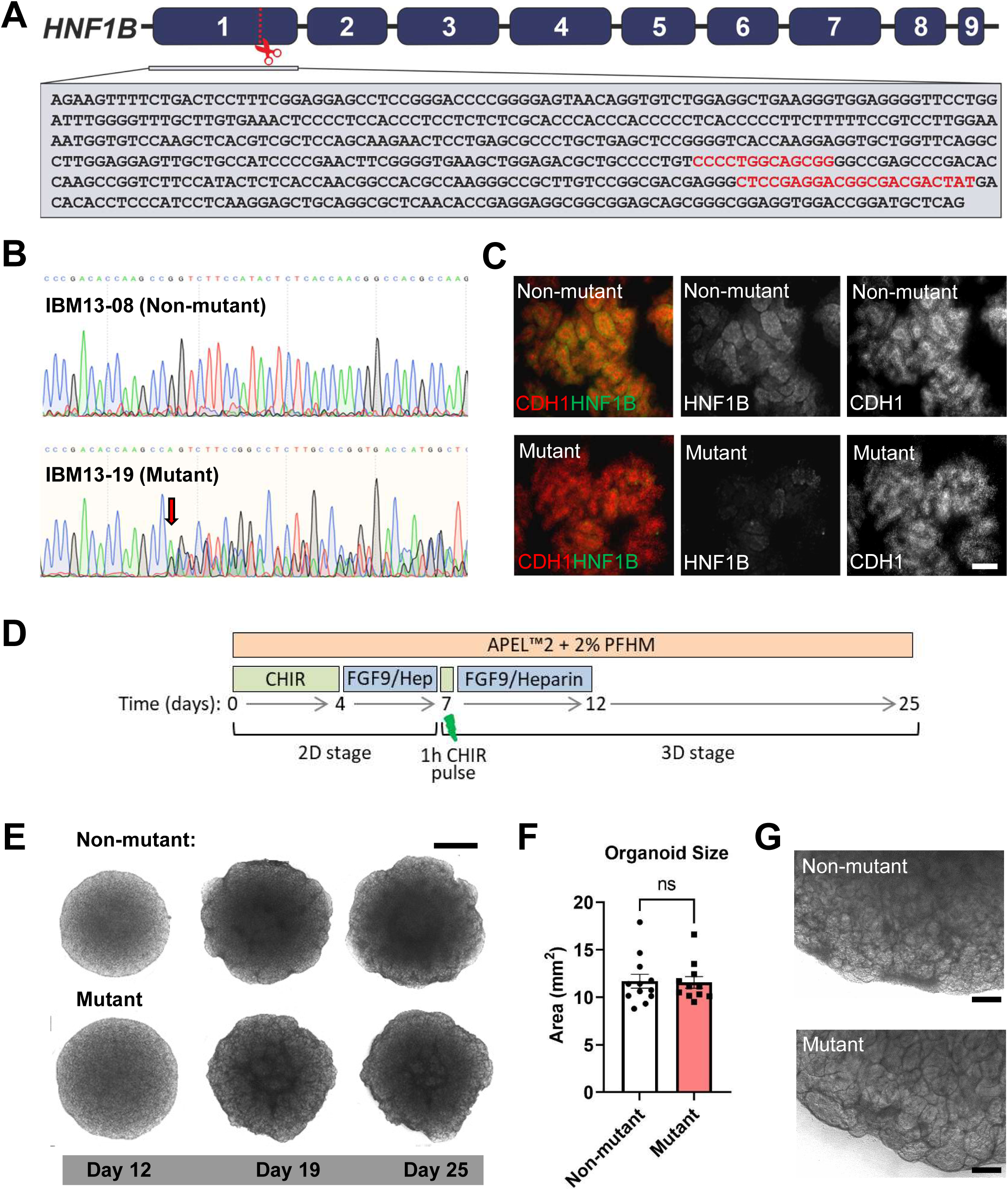
Generating heterozygous *HNF1B* mutant hESCs and their differentiation towards kidney tissues. **(A)** Sequence in exon 1 of *HNF1B*. Red dotted line and scissors mark the editing site, with gRNA binding sequences in red. **(B)** Sequencing chromatograms of exon 1 confirming the wild-type sequence in IBM13-08 (*non-mutant*) hESCs and the heterozygous frameshift starting (*red arrow*) at the CRISPR-targeted site in IBM13-19 (*mutant*) hESCs. **(C)** 2D kidney differentiation cultures (day 12) confirmed both lines formed CDH1+ aggregates but HNF1B immunoreactivity appeared less in mutant cells. **(D)** 25 day organoid protocol. **(E)** Phase contrast images of organoids. **(F)** Day 25 wild-type and mutant organoids had similar areas. **(G)** Mutant organoids contained bulkier internal structures than wild-type organoids. Bars: (C) and (G) 100 µM; (E) 1 mm.

**Figure 2.**
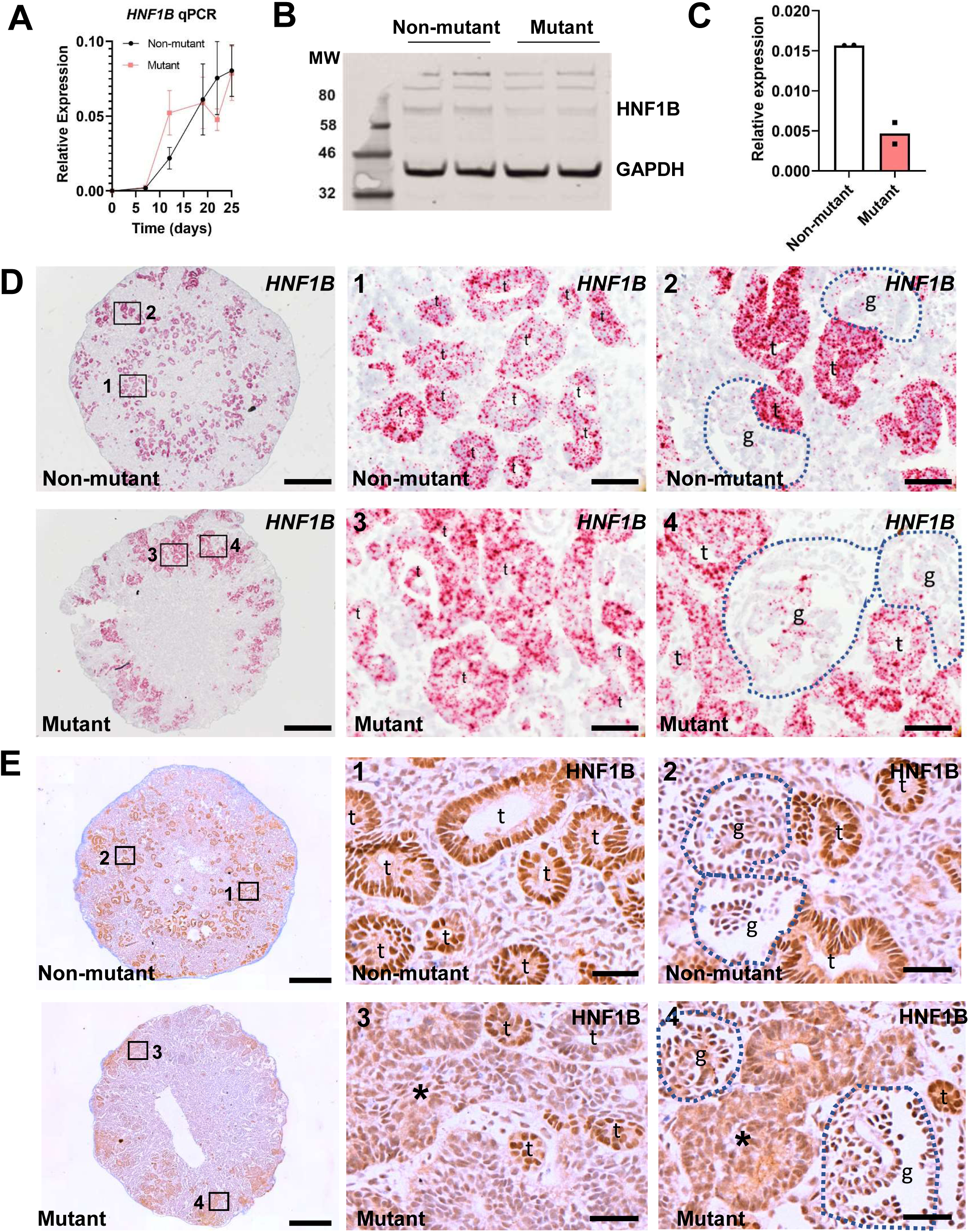
HNF1B in hESC-derived organoids. **(A)** RT-PCR using *HNF1B* exon 2 primers showed similar upregulation in *mutant* and isogenic *non-mutant* lines over 25 days. **(B)** Western blot of day 25 organoids (two organoid pools for each genotype) detected HNF1B doublets at 63 kDa. **(C)** Decreased HNF1B/GAPDH values in mutant organoids (average and individual values shown). **(D)** BaseScope ISH *HNF1B* (red dots) with nuclei counterstained blue with haematoxylin. Left-hand fames show low power overviews with enlarged areas depicted in boxes 1-4. Non-mutant tubules (*t* in 1) expressed *HNF1B* but there were barely any signals in glomeruli (*g* in 2). In mutants, *HNF1B* was expressed in bulky, morphologically aberrant, tubules (*t* in 3) and in tufts of aberrant looking glomeruli (*g* in 4). **(E)** HNF1B immunostaining (brown) in day 25 organoids, with haematoxylin counterstain. Left-hand frames show overviews with boxes 1-4 enlarged in the other frames. HNF1B was detected in nuclei of wild-type tubules (*t* in 1 and 2). In mutants, HNF1B was immunodetected in small calibre tubules (*t* in 3 and 4) but signals appeared attenuated and diffuse in bulky, morphologically aberrant, tubules (*asterisks* in 3 and 4). Bars: (D) 200 µM (overview) and 20 µM (enlargements); and (E) 500 µM (overview) and 40 µM (enlargements).

### Histology and functionality of hESC-derived organoids

Fig. 3A and B contrast histology overviews of control and mutant organoids. The mutant appeared to contain bulkier tubules that we proceeded to analyse in detail. In control organoids *Lotus tetragonolobus* lectin (LTL), a PT marker (Kishi et al., 2019), bound a subset of slender tubules in a uniform manner (Fig. 3C), whereas mutant dysmorphic tubules exhibited patchy staining (Fig. 3D). In native human kidneys, DTs and CDs are rich in CDH1 (Nouwen et al., 1993). In control organoids, a subset of tubules was CDH1+ (Fig. 3E), and mutant dysmorphic tubules showed patchy immunostaining (Fig. 3F). For both LTL+ and CDH1+ tubules, the cross sectional areas of mutant tubules were significantly larger than areas of wild-type tubules (Fig. 3K-M). *In vivo*, megalin and cubilin form a receptor complex on the apical (luminal) plasma membrane of PTs (Molitoris et al., 2022; Nielsen et al., 2016). Slender control organoid tubules displayed this pattern (Fig. 3C,G, I) but bulky mutant tubules showed predominantly cellular immunostaining of cubilin (Fig. 3H). Control organoids displayed an apical pattern for megalin (Fig. 3l) but this protein was undetectable in mutants (Fig. 3J). Synaptopodin is a podocyte actin-associated protein (Ning et al., 2020). Immunostaining (Fig. S1A and B) revealed synaptopodin+ glomeruli in control and mutant organoids but mutant glomeruli appeared larger with less compact podocyte tufts. Control and mutant organoids contained PECAM+ endothelia between tubules but neither genotype featured capillaries within glomerular tufts (Fig. S1C and D). We measured proliferation by detecting incorporated BrdU and apoptosis by activated caspase 3 immunostaining (Fig. S1E-J). BrDU+ tubule nuclei, and apoptotic figures were significantly increased across the whole organoid in mutants. Forskolin (FSK) is an adenylate cyclase activator that increases intracellular 3’,5’-cAMP. When intact metanephroi are exposed to FSK in organ culture, nephron lumina dilate, reflecting fluid transport and tubule functionality (Anders et al., 2013). Adding FSK to organoids (Fig. 4A) generated numerous translucent areas in controls whereas few were evident in mutants (Fig. 5B). Histologically (Fig. 4C-J), both the numbers of (Fig. 4C), and the area occupied by (Fig. 4D) these dilatations was significantly less in mutants. Fig. 4E-H depict histology overviews of organoids demonstrating no dilated structures in control (Fig. 4E) and mutant (Fig. 4F) organoids that were not exposed to FSK, and dilatations in FSK-exposed non-mutant (Fig. 4G) and infrequently in mutant (Fig. 4H) organoids. Probing with LTL or with CDH1 or synaptopodin antibodies (Fig. 4I-N) suggested that dilatations in non-mutant organoids affected PTs, DTs and glomeruli. The smaller dilated structures in mutants were harder to categorise, although some affected glomerular tufts (Fig. 4N).

**Figure 3.**
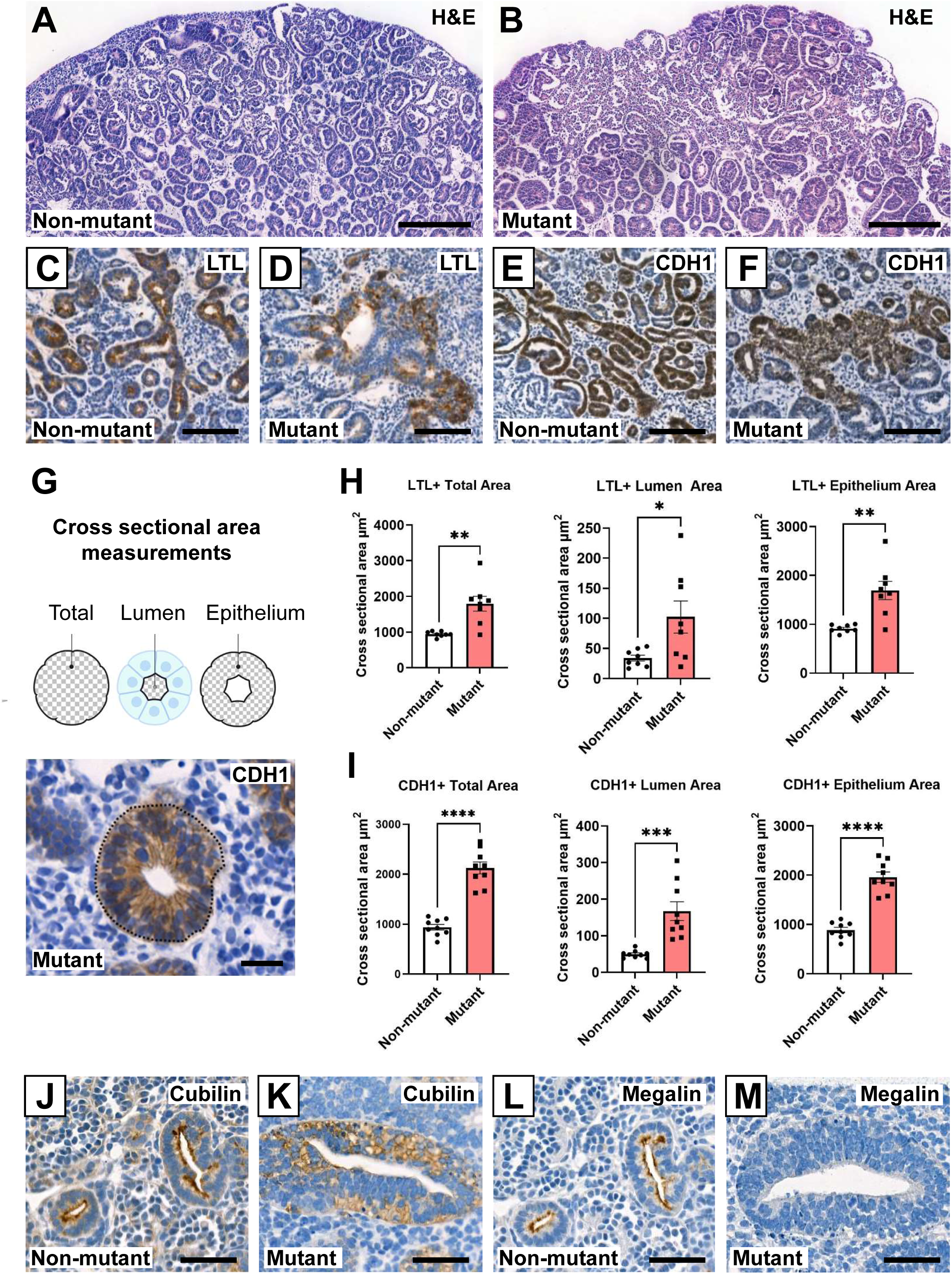
Aberrant tubules in *HNF1B* mutant organoids. Non-mutant **(A)** and mutant **(B** organoids counterstained with haematoxylin and eosin. Internal structures appeared larger in mutants. **(C)** Non-mutant and **(D)** mutant organoids stained with LTL (brown) with haematoxylin counterstain, showing slender LTL+ non-mutant tubules and bulky mutant tubules with patchy staining. **(E)** Non-mutant and **(F)** mutant organoids immunostained for CDH1 (brown) with haematoxylin counterstain, showing slender CDH1+ wild-type tubules and bulky mutant tubules with patchy staining. **(G)** Above: Cartoon showing the total, lumen and epithelium areas measured from perpendicularly cross-sectioned tubules. Below: Example tubule immunostained (brown) for CDH1. **(H)** Area of LTL+ profiles (mean±SEM; n=9 organoids from three independent differentiation experiments; *p<0.05, **p<0.005, t-test). **(I)** Areas of CDH1+ profiles (mean±SEM; n=9 organoids from three independent differentiation experiments; ***p<0.0005, ****p<0.00005 t-test). (**J** and **K**) Cubilin immunostaining (brown) showed an apical pattern in non-mutant tubules but a diffuse pattern in mutants. (**L** and **M**) Megalin immunostaining showed an apical pattern in non-mutant tubules (**L**) but megalin was not detected in mutant tubules. Bars: (A and B) 200 μM; (C-F) 100 μM; (G) 20 μM; and (J-M) 50 μM.

**Figure 4.**
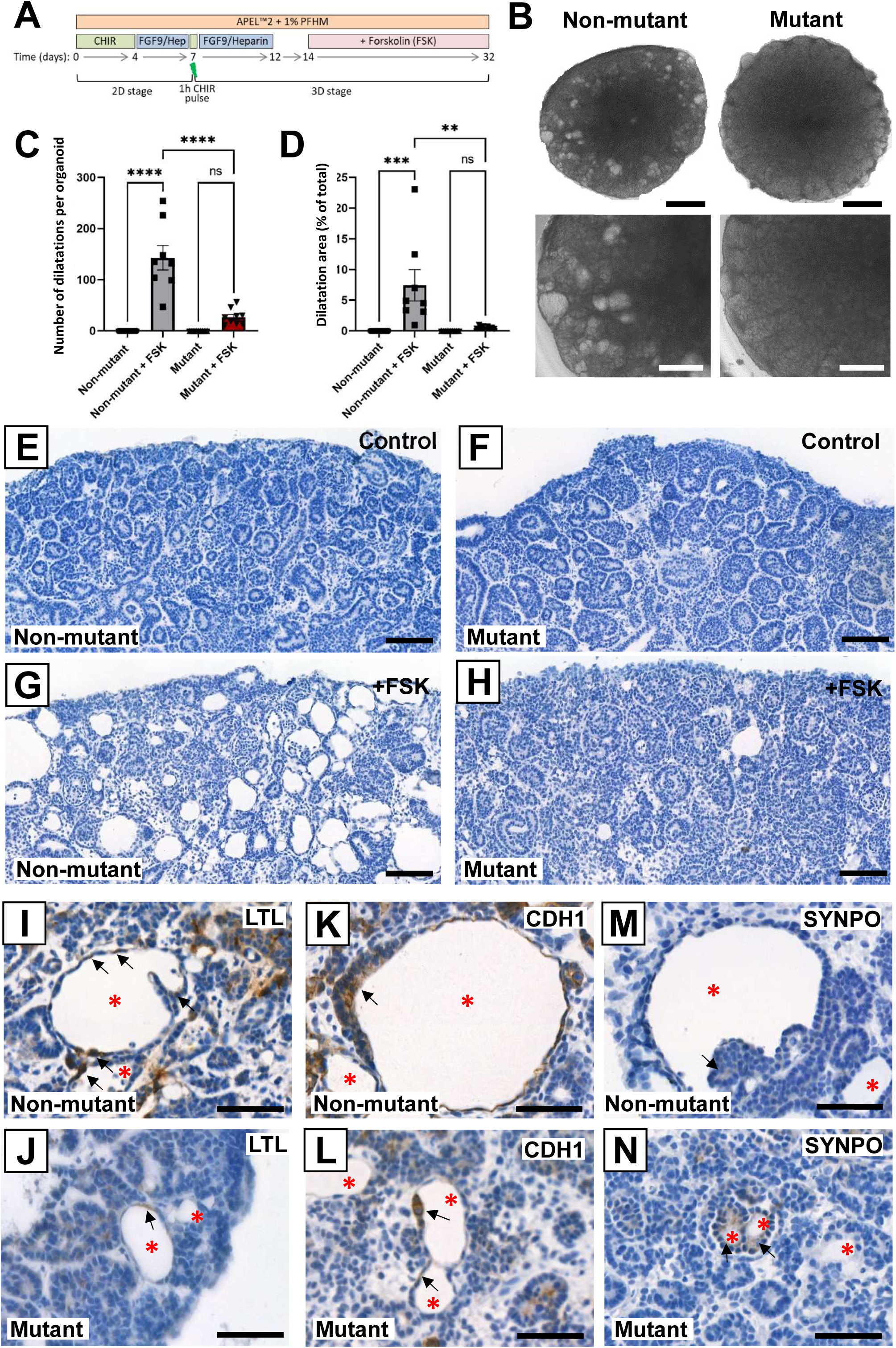
Deficient cAMP-induced lumen dilatation in *HNF1B* mutant organoids. **(A)** Organoids were exposed to forskolin (FSK) between 14 and 32 days. **(B)** Phase contrast images at day 32 showed that FSK had induced numerous dilated structures in non-mutant organoids but few in mutants. **(C)** Numbers of dilatations per organoid on histology (mean±SEM; n=9 organoids across 3 independent experiments; ****p<0.00005, one-way ANOVA with multiple comparisons). **(D)** Quantification of total dilated percentage area per organoid (mean±SEM; n=9 organoids across 3 independent experiments; ***p<0.0005, **p<0.005, one-way ANOVA with multiple comparisons). **(E-H)** Haematoxylin stained sections of non-mutant and mutant organoids, without (control) or with added FSK (+FSK). (**I to N).** Organoid sections reacted with LTL or immunoprobed for CDH1 or SYNPO and counterstained (blue) with haematoxylin. Red asterisks indicate dilated structures and black arrows indicate associated cells. Bars: (B) 1 mm (upper panels) and 500 μM (lower panels); (E-H) 200 μM; and (I-N) 50 μM.

**Figure 5.**
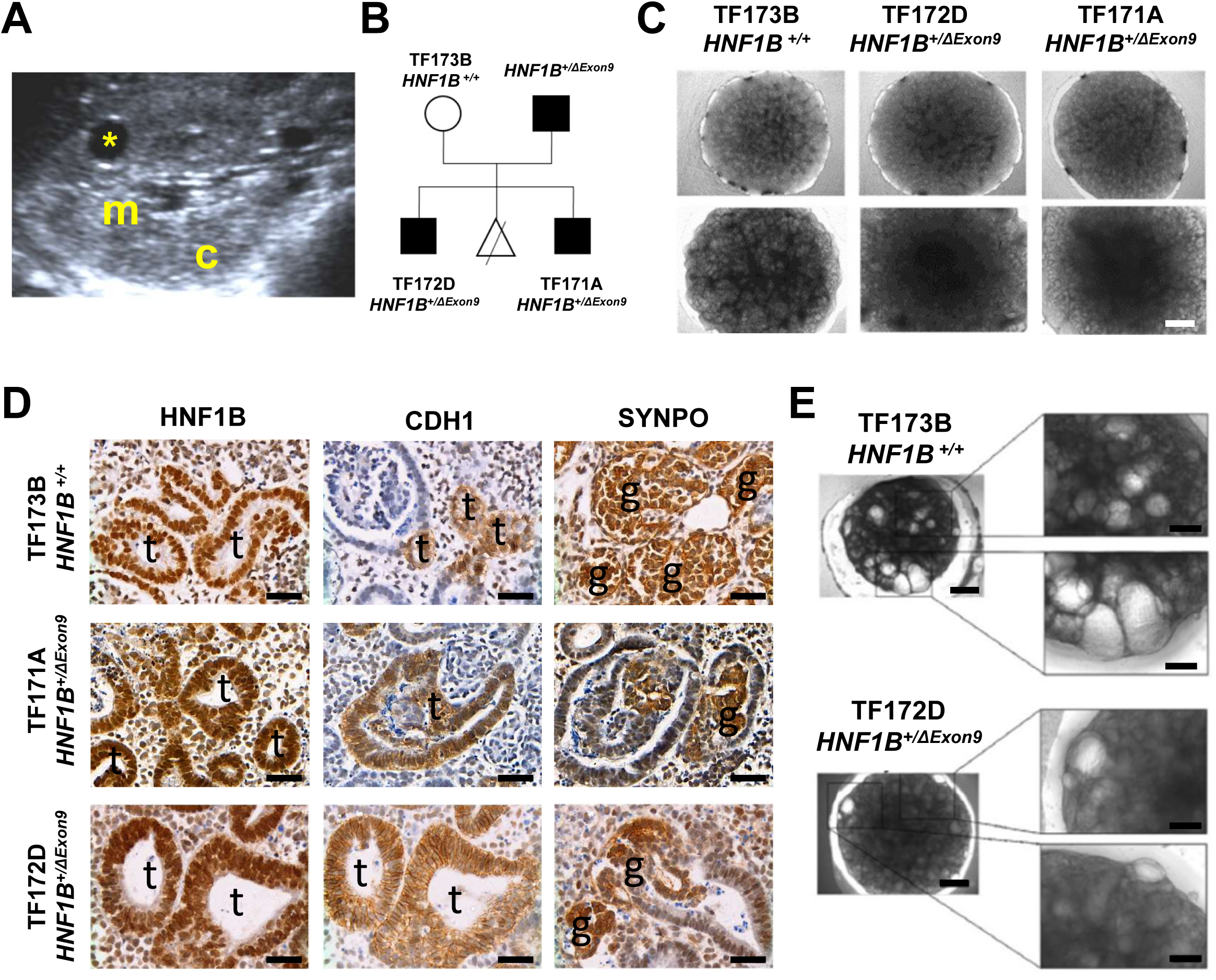
iPSC-derived organoids. (**A**) Kidney ultrasonography of affected *HNF1B*^+/ΔExon9^ male (TF172). Note lack of distinction between the echogenic cortex (c) and medulla (m); asterisk indicates a dilated structure. **(B)** Kindred from which iPSCs were derived. Black icons indicate individuals with DKMs. The triangle indicates the affected fetus who underwent elective termination. **(C)** Phase contrast images of organoids at day 15 (top images) and day 18 (bottom images). **(D)** Immunohistochemical comparison of *HNF1B^+/ΔExon9^*with *HNF1B^+/+^* control organoids. Note aberrant multi-layered CDH1+ and HNF1B+ tubules (t) and dysmorphic SYNPO+ glomeruli (g) in mutant tissues. Positive immunostaining (brown) with haematoxylin counterstain (blue). **(E)** 8-Br-cAMP-induced dilated structures appeared fewer in *HNF1B^+/ΔExon9^* organoids. Bars: (C) 500 μM; (D) 20 μM; (E) 500 µm (left frames) and 20 µm (right frames).

### Organoid differentiation from heterozygous *HNF1B* mutant patient-derived hiPSCs

To determine whether the phenotype of CRISPR/Cas9-mutated hESCs-derived organoids was representative of *HNF1B*-associated kidney disease, we evaluated organoids generated from hiPSCs derived from a family carrying an inherited heterozygous deletion of exon 9 (*HNF1B*^+/ΔExon9^) (Fig. 5A and B). hiPSCs from two affected brothers (TF171A and TF171D) and their clinically unaffected *HNF1B*^+/+^ mother (TF171B) were generated from PBMNs. The unrelated line SW162A was used as an additional non-mutant control. Genomic qPCR for exon 9 confirmed the heterozygous deletion in the brothers’ iPSCs (Fig. S2A). All hiPSC lines formed organoids (Fig. 5C). *HNF1B* transcripts, assessed using exon 2 primers, increased similarly during development in control and mutant organoids (Figure S2B). In contrast, using exon 9 primers, transcripts were lower in mutant organoids (Figure S2B). Dysmorphic CDH1+ tubules were detected in mutant organoids, contrasting with slender CDH1+ tubules in control iPSC-derived organoids (Fig. 5D). *HNF1B*^+/ΔExon9^ organoids contained malformed glomeruli, with Synaptopodin+ tufts surrounded by capsules comprised of bulky epithelia, in contrast to thin Bowman capsules in controls (Fig. 5D). We exposed organoids to 8-Br-cAMP that induces nephron dilatation in intact metanephroi maintained in organ culture (Anders et al., 2013). As for the FSK-exposed hESC-derived exon 1 mutant organoids, significantly fewer dilated nephrons were detected in 8-Br-cAMP-exposed *HNF1B*^+/ΔExon9^ organoids compared with control iPSCs-derived organoids (Fig. 5E; Fig. S2C and D).

### Overview of hESC-derived organoid transcriptomes

For model validity, we compared bulk RNAseq profiles of differentiating control hESCs with human kidneys at 10-12 weeks gestation when organs contain a nephrogenic cortex of MM and branching UB tips, with deeper layers of maturing nephrons, CDs and stromal cells (Woolf and Jenkins, 2015). Principal component analysis (PCA) showed that, as *in vitro* differentiation progressed over 25 days, organoid profiles approached those of native fetal kidneys (Fig. S3A). Heat maps (Fig. S3B-G) demonstrated that organoids expressed nephron precursor transcripts, and their levels tended to fall as organoids matured. Towards the end of the 25 day protocol, glomerular and PT transcript levels in organoids resembled those in fetal kidneys. In contrast, DT and certain CD transcripts did not generally reach fetal kidney levels. Transcriptional profiles of hESC-derived exon 1 heterozygous *HNF1B* mutant cells were compared with their control isogenic time-matched counterparts during differentiation. Assessed by PCA (Fig. S4A), the overall temporal progression was similar with samples clustering by day of development (0, 7, 12, 19 and 25), and controls and mutants clustered together. Levels of *HNF1B* in the RNAseq increased markedly from day 12, and similarly in the two genotypes (Fig. S4B). As organoid differentiation proceeded increasing numbers of significantly differentially expressed genes (DEGs) were identified between the controls and mutants (Fig. S4C): 107 at day 12; 151 on day 19; and 631 on day 25. Some transcripts differed at all three timepoints, whereas others differed on only one or two of these days (Fig. S4D). The RNAseq was interrogated for DEGs containing the canonical *HNF1B* binding motif upstream of their transcription start site (Adalat et al., 2009) (Fig. S5A). This identified increasing numbers of such genes on days 12, 19 and 25 (Fig. S5B-E) and, while similar numbers of genes were up- or downregulated on days 12 and 19, most DEGs on day 25 were downregulated. Gene ontology (GO) enrichment analysis was undertaken, comparing mutant and isogenic control organoids for biological process (Fig. S6A-C), molecular function (Fig. S6D-F) and cellular compartment (Fig. S6G-I) enrichment. Among the deregulated terms, calcium ion binding, cadherin binding, collagen containing extracellular matrix, and integral component of plasma membrane were prominent.

### Characteristic kidney transcripts in *HNF1B* mutant hESC-derived organoids

We examined organoids for transcripts known to be specific for, or enriched in, certain cell types in native kidneys, with results depicted in heat maps (Fig. 6A-H) and volcano plots (Fig. 6I-K). Transcripts encoding glomerular basement membrane molecules (Fig. 6A) implicated in genetic diseases (Boyer et al., 2017) (*COL4A3, COL4A4, COL4A5, LAMB2*) were expressed similarly between genotypes, while *LAMB1* was significantly lower in day 25 mutants (Fig. 6K). Glomerular podocyte genes *NPHS1, NPHS2, PODXL* and *SYNPO1* became upregulated during organoid culture, with a significantly higher level of *PODXL* in day 19 mutants (Fig. 6K). WT1 is expressed in the nephron lineage, with highest levels in podocytes (Winyard et al., 1996b), and was detected at day 12, increasing similarly in controls and mutants.

**Figure 6.**
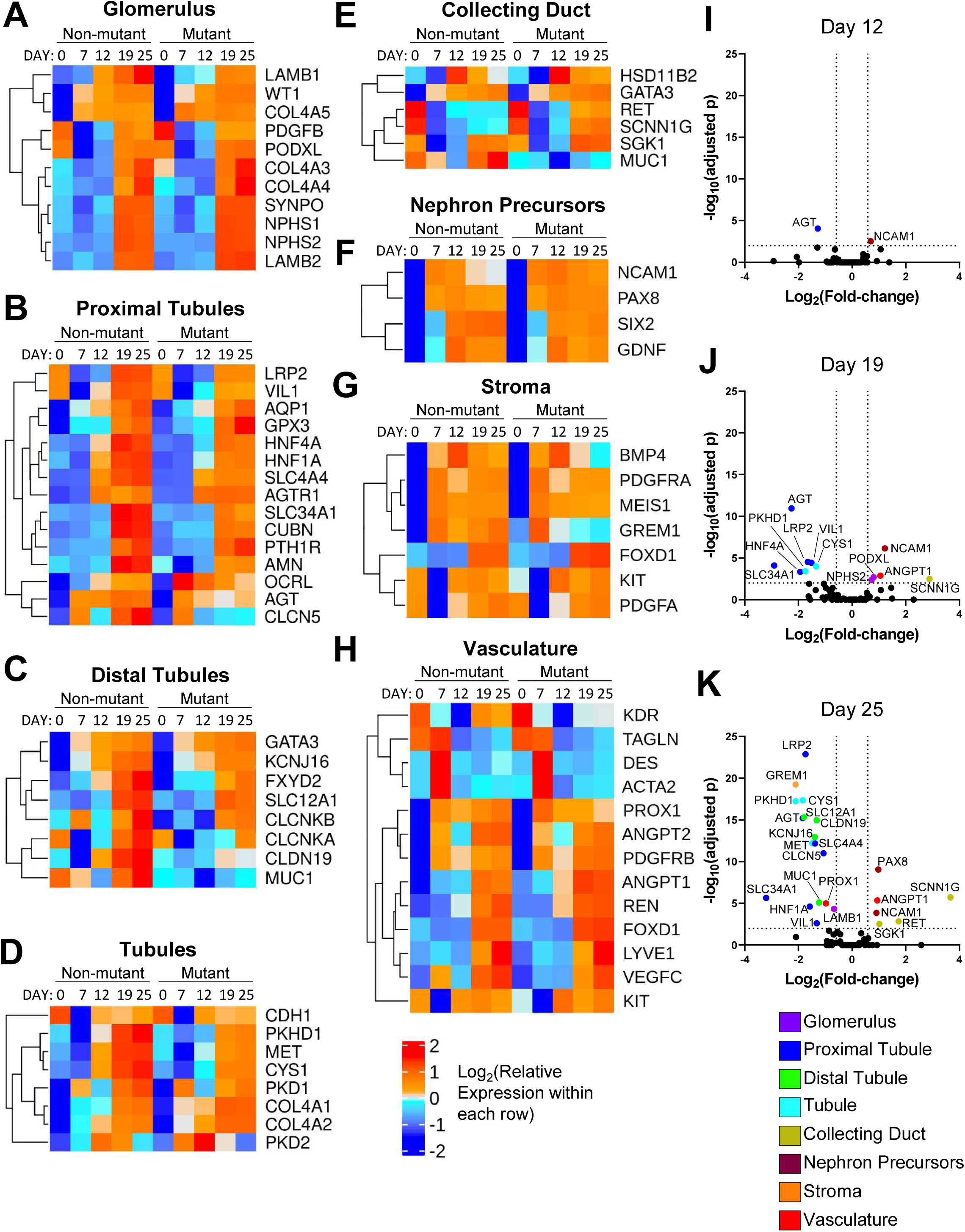
Profiles of established kidney transcripts in hESC-derived organoids. **(A-H)** Heatmaps through the differentiation protocol, with days 12, 19 and 25 being the organoid phase. Each element in the heat maps represents the mean of three independent differentiation experiments. Genes were included if they had an average read count>50 on at least one day of organoid differentiation. **(I-K)** Volcano plots showing significantly deregulated transcripts at days 12 (I), 19 (J) and 25 (K) with cut-offs of a log2 fold change of 0.5 and log10 p-adjusted significance value of 2. See main Results text for detailed interpretation.

Strikingly, several genes implicated in Mendelian PT diseases (OMIM) showed deregulated expression in mutant organoids (Fig. 6B, J and K). *LRP2*, encoding Megalin and mutated in Donnai-Barrow syndrome (OMIM 222448), was expressed at low levels on day 12 and was then upregulated but with significantly lower levels in day 19 and 25 mutants (Fig. 6J and K). *CLCN5,* the chloride channel mutated in Dent disease 1 (OMIM 300009), was detected in organoids but with significantly lower levels in day 25 mutants (Fig. 6K). *SLC34A1* encodes a brush-border Na^+^-phosphate cotransporter mutated in hypophosphataemia (OMIM 612286) and renal Fanconi syndrome 2 (OMIM 613388). It was not detected at day 12 and was then upregulated but with significantly lower levels in day 19 and 25 mutant organoids (Fig. 6J and K). *SLC4A4* encodes a sodium/bicarbonate cotransporter mutated in PT acidosis (OMIM 604278). It became upregulated in organoids but with significantly lower levels in day 25 mutants (Fig. 6K). *HNF1A*, a transcription factor binding partner of HNF1B causing PT dysfunction when mutated in mice (Pontoglio et al., 1996), was upregulated during organoid culture, with significantly lower levels in day 25 mutants (Fig. 6K). BaseScope for *HNF1A* (Fig. S7) detected transcripts in control organoid tubules, will a lower signal in mutants. *HNF4A,* mutated in Fanconi renotubular syndrome, was upregulated in organoids but significantly less in day 19 and 25 mutant organoids (Fig. 6J and K). *AGT*, encoding the angiotensin precursor and mutated in renal tubular dysgenesis (OMIM 267430), was detected but with significantly lower levels through the mutant organoid phase (Fig. 6I-K). *VIL1,* encoding brush border Villin, was upregulated through differentiation but with significantly lower levels in day 19 and 25 mutants (Fig. 6J and K). These aberrations could not simply be attributed to a global failure of PT differentiation because other characteristic PT transcripts were expressed similarly in mutant and control organoids: *AMN* and *CUBN*, encoding proteins forming a receptor complex with Megalin; *PTH1R*, encoding parathyroid hormone receptor; *AQP1* encoding a water transporter; and *GPX3* and *OCRL* encoding tubular enzymes.

Certain characteristic DT transcripts (Fig. 6C) were aberrantly expressed in mutant organoids. *CLDN19*, mutated in hypomagnesemia type 5 (OMIM 248190), was expressed by organoids but at significantly lower levels in day 25 mutants (Fig. 6K). Similar aberrant patterns (Fig. 6K) were found for: *MUC1*, encoding a mucin mutated in medullary cystic kidney disease (OMIM 174000); *SLC12A1*, encoding the Na^+^-K^+^-Cl^-^ transporter mutated in Bartter syndrome type 1 (OMIM 601678); and the potassium channel *KCNJ16* mutated in hypokalaemic nephropathy and deafness (OMIM 619406). In contrast, *CLCNKB*, a chloride channel, and *GATA3*, a transcription factor expressed in DTs and CDs, were expressed similarly in controls and mutants. *UMOD*, coding for uromodulin, was not detected in organoids. Other genes, widely expressed in kidney tubule epithelia, were downregulated in mutants (Fig. 6D and I-K) including: *MET* (Kolatsi-Joannou et al., 1997), encoding the HGF receptor, at day 25; autosomal recessive PKD associated gene *CYS1*, encoding cystin (Yang et al., 2021) at days 19 and 25, and *PKHD1*, encoding Fibrocystin (Nakanishi et al., 2000), at days 19 and 25. In contrast, *PKD1* and *PKD2*, coding for the polycystins mutated in autosomal dominant PKD (Chauvet et al., 2002), were expressed similarly in controls and mutants, as was *CDH1*, *COL4A1* and *COL4A2*. Regarding CD genes (Fig. 6E, J and K), *AVPR2*, encoding a vasopressin receptor, and the water channel *AQP2* were not expressed in organoids. *SCNN1G* encodes a CD sodium channel mutated in Liddle syndrome (OMIM 618114) and pseudohyoaldosteronism type 1 (OMIM 264350); it was not detected in controls but was expressed in day 25 mutant organoids. Similar upregulation was noted for *SGK1* encoding serum/glucocorticoid-regulated kinase 1, and for *RET* encoding a tyrosine kinase receptor expressed by branching UB tips in native metanephroi. In contrast, *HSD11B2*, encoding 11-beta-hydroxysteroid dehydrogenase type II, was expressed similarly in the two genotypes.

We examined transcripts expressed by MM and primitive nephrons (Fig.6F). Compared with wild-type organoids, the cell-cell adhesion molecule *NCAM1* was expressed at higher levels in mutant orgnoids at days 12, 19 and 25 (Fig. 6I-K). *PAX8* was upregulated in mutant organoids at day 25 (Fig. 6K). In contrast, *GDNF*, a RET ligand, and the *SIX2* transcription factor were expressed similarly in the two genotypes. Kidney interstitial stromal cell markers (Wilson et al., 2021; Bantounas et al., 2021) *FOXD1*, *KIT*, *MEIS1*, *PDGFA* and its receptor *PDGFRA*, were detected in organoids, with no difference between genotypes (Fig. 6G) *GREM1*, encoding a protein antagonising BMP4, which itself inhibits UB branching (Michos et al., 2007), was significantly lower in day 25 mutant organoids (Fig. 6K). We investigated transcripts implicated in differentiation of endothelia and vascular smooth muscle cells (SMCs) (Gnudi et al., 2015; Jafree et al., 2019) (Fig.6H). *ANGPT1* was expressed at significantly higher levels in mutant organoids on days 19 and 25 (Fig.6J and K), whereas its antagonist *ANGPT2* was expressed similarly in the two genotypes. *VEGFA, VEGFC* and *PECAM1* were expressed similarly in control and mutant organoids. *PROX1* and *LYVE1*, lymphatic endothelial genes, were expressed in organoids, with a significant decrease in *PROX1* in day 25 mutants (Fig. 6K). *ACTA2*, *DES*, and *TAGLN* encoding SMC cytoskeletal proteins, *REN* expressed in juxtaglomerular SMCs, and *PDGFRB* expressed in mesangial cell precursors, were detected at similar levels in control and mutant organoids.

### Deregulated glutamate receptor and pathway genes in mutant organoids

*GRIK3*, encoding glutamate ionotropic receptor (iGluR) kainate type subunit 3 (Hansen et al., 2021), was among the ten most deregulated transcripts in mutant organoids. *GRIK3* levels in human fetal kidneys were similar to those in control organoids (Fig. S8A) but *GRIK3* was significantly upregulated on days 12, 19 and 25 in mutant compared with control organoids (Fig. S8B). On the final day of culture, *GRIK3* expression was 10 times higher (Padj=1.95E-31) in heterozygous *HNF1B* mutant than in control organoids (Fig.7A). Western blotting of organoid protein showed significantly increased GRIK3 protein in mutant cultures (Fig. 7B and C). BaseScope ISH (Fig. 7D) of control organoids detected *GRIK3* transcripts in tubules and more sparsely in interstitial cells, while in mutant organoids *GRIK3* was prominently expressed in dysplastic tubules. Immunohistochemistry for GRIK3 showed a predominantly tubular pattern in both control and mutant organoids (Fig. 7E). We next undertook GRIK3 immunostaining on sections of third trimester human fetal kidneys (Fig. S9). GRIK3 was detected in tubules in a control third trimester kidney; some such tubules had apical staining with periodic acid Schiff (PAS), marking them as PTs. In a section from a fetus carrying a heterozygous *HNF1B* mutation, GRIK3 was prominent in dysplastic tubules, some also likely PTs as apically stained with PAS.

**Figure 7.**
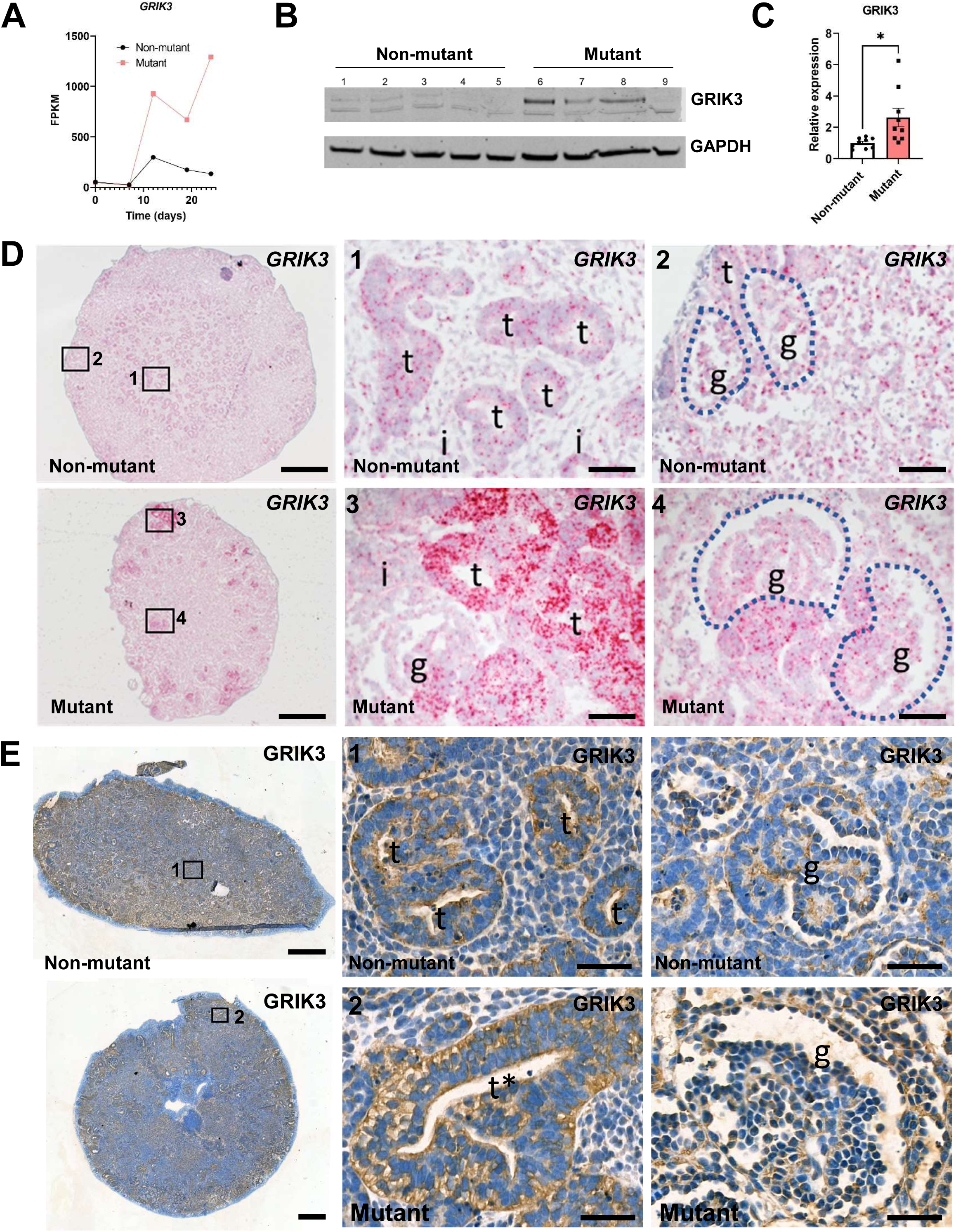
GRIK3 in kidney organoids. **(A)** RNAseq average reads for *GRIK3* during differentiation of non-mutant and *HNF1B* mutant hESCs, showing increased levels in mutants organoids (days 12, 19 and 25). **(B)** GRIK3 Western blot with 5 non-mutant and 4 mutant samples. **(C)** Quantification confirmed increased GRIK3/GAPDH in mutant organoids (mean±SEM; n=9, across four independent differentiation experiments; *p<0.05, t-test). **(D)** BaseScope for *GRIK3* with signals appearing as red dots; nuclei were counterstained blue with haematoxylin. Left-hand images are low power overviews of day 25 organoids, and other frames show high power images (1-4). In non-mutant organoids *GRIK3* was expressed in tubules (*t*) with sparser signals in interstitial cells (*i* in 1) and glomeruli (*g* in 2). In mutant organoids, *GRIK3* was highly expressed in large dysmorphic tubules (*t* in 3), with transcripts also noted in aberrant glomeruli (*g* in 4). **(E)** GRIK3 immunostaining (brown). Left-hand images are overviews of day 25 organoids, and the other frames (1-4) are high power images. In non-mutant organoids GRIK3 was immunodetected in tubules (*t* in 1). In mutant organoids, GRIK3 was prominent in multi-layered dysplastic tubules (*t* and *asterisk* in 3). A low level of immunostaining was noted in glomeruli (*g*) of both genotypes. Bars: (D) 200 μM (left frames) and 20 μM (other frames); (E) 500 μM (left frames) and 40 μM (other frames).

Kyoto encyclopaedia of genes and genomes (KEGG) enrichment analysis indicated that ‘glutamatergic synapse signalling’ pathways differed between mutant and control organoids (Fig. S10A; Table S4). We therefore focussed on transcripts encoding other ionotropic glutamate receptor (iGluR) subunits (Hansen et al., 2021; Valdivielso et al., 2020). Genes encoding several subunits belonging to the kainate (KA) and N-methyl-D-aspartate (NMDA) receptor subfamilies were expressed in control and mutant organoids (Fig. S8A-E). *GRIN2B* was significantly increased in day 12 mutant organoids, and *GRID1* was significantly increased at 19 and 25 days in mutants. Conversely, *GRIN3A* was significantly increased in day 19 mutant organoids, while *GRIN2A*, *GRIK5,* and *GRIA3* were expressed in organoids at similar levels between genotypes. *GRIN1* was not expressed in organoids and was only expressed at a low level in fetal kidneys. We examined expression of other genes participating in glutamate signalling (Hansen et al., 2021) but which do not code for iGluRs (Fig. S8F). For glutamate transporters, *SLC7A11* was upregulated in day 12 mutant organoids, while *SLC1A1* was downregulated in day 25 mutants. *HOMER2* encodes a scaffold protein binding various Ca^2+^ conducting channels (Jardin et al., 2013) and was upregulated in day 19 and 25 mutant organoids, and *CAMK2A* (Stephenson et al., 2017), a mediator of intracellular Ca2+ signalling and a participant in the NMDA receptor signalling complex, was upregulated in day 25 mutant organoids. *SYT1* coding for Synaptotagmin1 (Rastaldi et al., 2006) was upregulated in day 25 mutants. *PLCB1,* encoding a downstream mediator of glutamate signalling (Lo Vasco et al., 2004), was downregulated in mutant organoids at day 25, as was *SHANK2*, encoding a scaffold protein interacting with iGluRs (Schmeisser et al., 2012). Supporting the prevalence of transporter deregulation, it was notable that GO Cellular compartment enrichment analysis revealed significant gene enrichment in terms associated with the cell surface (Padj<0.01, Fig. S6 G-I) notably well over 100 genes in the term ‘Integral components of plasma membrane’ at d12 19 and 25.These included *LRP2, PODXL, NPHS2, PKHD1* and *MUC1* as well as 11 members of the extensive membrane transporter SLC genes, across a number of solute transport subfamilies, at day 19 and 25.

### Other pathways deregulated in *HNF1B* mutant organoids

Given that glutamate signalling impacts on intracellular calcium levels, it was notable that KEGG analysis (Table S4) indicated several differences in transcripts in calcium signalling pathways between mutant and control organoids (Fig. S10B). KEGG analysis also suggested that several components of WNT signalling pathways were dysregulated in mutant organoids (Fig. S10C). WNT genes are essential for several steps in kidney development (Halt et al., 2014). The following WNT pathway genes were decreased in mutant organoids: the ligands *WNT5A* (day 12), and *WNT10B* (day 25); *FZD4*, encoding a WNT receptor implicated in both the canonical and a non-canonical planar cell polarity pathway (day 25); and *DKK1* encoding a WNT pathway inhibitor (days 19 and 25).

## DISCUSSION

Our results reveal important morphological, molecular, and physiological roles for HNF1B in human kidney tubule morphogenesis and functional differentiation and suggest druggable targets to ameliorate disease.

### Heterozygous *HNF1B* mutant organoids mimic certain features of human *HNF1B*-associated DKMs

Human DKMs initiate organogenesis, yet their internal tissue organisation is deranged (Winyard et al. 1996a and 1996b; Kohl et al., 2022). With regard to *HNF1B*-associated DKMs, histological features include multi-layered large tubules and dysmorphic glomeruli, sometimes with dilated Bowman spaces (Bingham et al., 2002; Humaitre et al., 2006; Duval et al., 2016; Nakayama et al., 2022). Given that dysplastic tubules do not exactly resemble any normal structures in the developing or mature kidney, their nephron segment or CD origin has been unclear. In our study, we used a hPSC differentiation protocol which generates organoids that are rich in nephron components, especially glomeruli and PTs, but which do not contain differentiated CDs (Bantounas et al., 2018 and 2021; Takasato et al., 2015; Wu et al., 2018; Howden et al., 2021). Indeed, this pattern was supported by comparison of RNAseq transcript expression between control organoids with first trimester human fetal kidneys. Detailed histological examination of non-mutant organoids showed that they contained avascular glomeruli together with tubules that bound LTL, consistent with a PT identity (Kishi et al., 2019), or which reacted with CHD1 antibody, consistent with a DT identity (Nouwen et al., 1993). Our study showed that heterozygous mutant *HNF1B* hESCs or hiPSCs can form tissues containing kidney-like structures within organoids. This is consistent with the great majority of clinical reports noting that individuals with *HNF1B* mutations do have kidneys, in sharp contrast to certain other human genetic diseases (e.g. *FRAS1* or *FREM2* mutations) where organogenesis fails to initiate and the phenotype is an absent organ (kidney agenesis) rather than dysplasia (Woolf, 2022).

### Roles for *HNF1B* in human kidney tubule morphogenesis and functional differentiation

The main abnormal morphological feature of heterozygous *HNF1B* mutant organoids was a significant increase of tubule diameter, with some tubules being multi-layered rather than having a normal simple single layered epithelial wall. Significantly, this was found in both CRISPR-mutant hESC-organoids and those derived from reprogrammed iPSCs from affected individuals. Mutant organoids displayed lower expression than non-mutants of numerous transcripts characteristic of PTs and DTs. These included certain genes (e.g. *CLCN5, SLC34A1, SLC4A4, CLCNKA* and *SLC12A1*) implicated in Mendelian diseases (OMIM) where tubules fail to functionally mature, resulting in urinary wasting of electrolytes and low molecular weight proteins, together with acid-base aberrations. Indeed, deficiencies of these tubular transcripts helps to explain the clinical phenotype of urinary electrolyte wasting reported in *HNF1B*-assocated DKMs (Adalat et al. 2002 and 2019). A trivial explanation for these molecular results would be that no tubules with PT- or DT-like identities form in mutant organoids. This hypothesis is not supported by the fact that several other genes characteristic of these nephron segments were similarly expressed in mutant and non-mutant organoids. Another explanation could be that tubule maturation is simply delayed in mutant organoids. This is difficult to definitively refute because the *in vitro* protocol does not allow viable tissue maintenance beyond several weeks. We, however, favour another hypothesis, namely that heterozygous *HNF1B* mutations lead to the generation of aberrant epithelia with features not found in regular non-mutant kidney tissues. These features include enlarged and hyperproliferative tubules that have hybrid identities including binding to LTL and expressing CDH1, and disordered polarity as evidenced by diffuse cytoplasmic immunostaining for cubilin rather than the normal apical membrane pattern. This represents an abnormal cell fate rather than incomplete normal differentiation. Our molecular analyses also suggest that *HNF1B* mutant organoids display a modest upregulation of UB/CD lineage genes such as *RET, SGK1* and *SCNN1G* and this is consistent with metaplastic shift of cell identity within mutant organoids. In fact, it has been reported that nephron-rich hPSC-kidney organoids display a degree of plasticity such that by manipulating them biochemically they can be redirected to a UB/CD identity (<u>Howden et al., 2021</u>). Our results with heterozygous mutant hESCs and hiPSCs generally support those of a preliminary analysis of homozygous *HNF1B* mutant organoids reporting few LTL+ tubules compared with wild-type organoid (Przepiorski et al, 2018).

### Heterozygous *HNF1B* mutant organoids do not form cysts

A radiological feature of *HNF1B*-associated DKMs are cysts, up to a few centimetres across, and, on histology, glomeruli have dilated Bowman spaces and tubules have dilated lumina (Humaitre et al., 2006; Duval et al., 2016; Nakayama et al., 2022). In contrast, although mutant organoid tubules contained modestly but statistically significantly larger lumina than those of wild-type tubules, overt cysts were not present under standard development conditions. This is despite the facts that heterozygous *HNF1B* mutant organoids showed marked downregulation of certain genes (e.g., *CYS1* and *PKHD1*) that maintain a healthy non-cystic phenotype in kidney epithelia (Yang et al., 2021; Nakanishi et al., 2000). Our organoids lacked features of CDs, so it is possible that, had these been present, cysts may have occurred. Of note, another group generated heterozygous *HNF1B* mutant hPSC and differentiated them into UB/CD-like cells (Mae et al., 2020). These cells formed UB/CD organoids and in mutants “the number of budding regions tends to be reduced” but cysts were not observed (Mae et al., 2020). In certain genetic human cystic kidney diseases, such as those associated with mutations in *PKD1* or *PKD2*, *in vivo* kidney cystogenesis is largely driven by cAMP signalling (Richards et al., 2021). Compared with control counterparts, HNF1B mutant organoids resisted chemical induction by cAMP which generated dilated nephrons in control organoids. This resistance is consistent with a lack of mature function of mutant tubules, in accord with the lower expression of many tubular transporters evidenced by the RNAseq. We conclude that the genesis of cysts in *HNF1B*-associated DKMs may be a secondary, and late, feature in the development of the phenotype, and not driven by cAMP signalling. Perhaps the presence of glomerular filtration in conjunction with a primary aberration in tubule biology are needed to generate overt cysts in *HNF1B* disease. In future, implantation of heterozygous *HNF1B* mutant organoids into mice, as described for non-mutant hPSC-kidney progenitors (Bantounas et al., 2018; Bantounas et al., 2020), may allow prolonged growth of mutant tissues together with vascularisation of glomeruli and generation of ultrafiltrate. In this setting, dilatation may then occur in glomeruli and tubules. Compared with the morphological and molecular derangements of tubules in heterozygous *HNF1B* mutant organoids, mutant glomeruli had a milder phenotype, appearing to have more prominent tufts with generally preserved expression of podocyte genes. This is consistent with the normal expression pattern of *HNF1B* in native human kidneys (Kolatsi-Joannou et al., 2001) being confined to the tubule epithelia.

### Molecular mechanisms of *HNF1B*-associated DKMs

Our RNAseq analyses of mutant organoids identified the deregulation of many genes containing the canonical HNF binding sequence (Adalat et al, 2009). Several were genes characteristically expressed by kidney tubule epithelia including *CLCN5, CYS1, HNF1A, HNF4A, KCNJ16, MUC1, PKHD1* and *SLC34A1.*Thus, it is likely that their deregulated expression is a direct effect of decreased functional HNF1B protein. On the other hand, several key genes involved in kidney development, such WNT pathway transcripts, were also deregulated yet do not contain the canonical HNF1B binding site. These changes may be secondary to direct HNF1B induced gene regulation, yet their altered expression may contribute to the DKM-like phenotype. One of the most prominent examples of such genes was *GRIK3,* encoding a glutamate receptor, that was markedly upregulated through the organoid phase of heterozygous *HNF1B* mutant differentiation, and to a much greater extent than in controls. In support of a role for glutamate receptors in disease, KEGG pathway analysis predicted deregulation of glutamatergic signalling. Further, histological analyses of *GRIK3* transcripts and protein revealed expression predominantly in tubule epithelia in non-mutant organoids, with noticeably higher expression in dysmorphic tubules in *HNF1B* mutant organoids. Similar differential expression between mutant and control kidney was seen in fetal kidney sections. Moreover, several other genes that code for glutamate receptors or transporters, or components of glutamatergic intracellular signalling machinery, were aberrantly expressed in mutant organoids. These observations are highly novel, especially given that glutamate receptors have hitherto not been studied in either normal or abnormal human kidney development.

Glutamatergic signalling and glutamate metabolism have been intensively studied in neuronal tissues where glutamate acts as an excitatory neurotransmitter (Hansen et al., 2021). But roles for glutamatergic signalling, and GRIK3 itself, are emerging outside the nervous system. GRIK3 expression is prominent in breast cancer tissues and *in vitro* it increases proliferation and migration of breast cancer cells, driving epithelial to mesenchymal transition (Xiao et al., 2019). Similarly, GRIK3 has been implicated in proliferation and migration of intestinal and lung cancer cells (Du et al., 2020). Glutamic acid impairs mouse blastocyst development and glutamate receptors, inlcuding GRIK3, are involved in this effect (Spirkova et al., 2022). Of note, components of the glutamate signalling system have been identified in mature mammalian kidneys *in vivo* and in kidney cell epithelial cell lines (Hediger, 1999; Valdivielso et al., 2020) e.g. *GRIN2B* in hypoxic murine kidney tubules (Iwata et al., 2022) and *GRIN2A* in murine kidney tubules (Iwata et al., 2022) *with SLC7A11*, and SLC1A in PT cells (Wang et al., 2020; Shayakul et al., 1997; Welbourne et al., 1999). Importantly, increased monosodium glutamate dietary intake in rats leads to increased glomerular filtration rate, which the NMDA receptor antagonist MK-801 reduced (Mahieu et al., 2016). Glutamate transporters move extracellular glutamate into PTs, facilitating glutamine/glutamate metabolism, urinary acidification and movement of bicarbonate back into the body (Welbourne et al., 1999). Knockdown of GRIN1 (NMDAR1) in a PT cell line led to an epithelial to mesenchymal transition as evidenced by changed morphology and upregulation of α-smooth muscle actIn, while addition of NMDA blunted *in vitro* expression of mesenchymal markers stimulated by TGFB1 (Bozic et al., 2011). Moreover, *in vivo* treatment with NMDA ameliorated kidney fibrosis triggered by experimental ureteric obstruction (Bozic et al., 2011). In mice receiving chemical NMDAR blockade, urinary protein levels rise while, *in vitro*, antagonising NMDAR causes cytoskeletal remodelling in podocytes (Giardino et al., 2009). These diverse observations indicate that glutamate and glutamatergic signalling impact on the health of kidney epithelial cells in mature organs. They are also consistent with the hypothesis that the deregulation of numerous glutamatergic signalling genes such as *GRIK3* play roles in the pathobiology of kidney dysplasia associated with *HNF1B* mutation.

An unanswered question is whether developing kidneys are exposed to glutamine/glutamate *in vivo*. Nevertheless, free amino acids are present in the milieu of early developing embryos (Van Winkle., 2021) and human cord blood at term contains glutamate, with levels increased in the presence of fetal distress (Perez-Mato et al., 2016). In the central nervous system, astrocytes synthesize and release glutamine that is taken up by neurons to generate glutamate to be used in neurotransmission (Anderson and Schousboe, 2022). Whether a similar glutamate-producing mechanism operates in the kidney is not known. Alternatively – or in addition to glutamate – kidney GluRs may function to sense other amino acids in the proto-urine: For example, D-Serine binds to and activates NMDARs in the kidney resulting in Ca^2+^-mediated increase in reactive oxygen species, which leads to renal insufficiency in mice. Many of the resulting symptoms are reversible by the use of NMDAR inhibitors (Tseng et al, 2021). Finally, we note that drugs that modulate glutamate signalling are being explored as treatments in non-renal (e.g. brain) diseases (Stone, 2011) and this in turn suggest that similar drugs may ameliorate features of kidney organogenesis associated with *HNF1B* mutations.

## METHODS

### Sources of human tissues

First trimester fetal kidneys collected after maternal consent and ethical approval (ethics REC 08/H0906/21+5; and REC 18/NE/0290) and were provided by the MRC and Wellcome Trust Human Developmental Biology Resource (http://www.hdbr.org/). Third trimester human fetal kidneys, were used for research histology studies with approval from the Ethics committee of the Hôpital Robert Debré, as detailed previously (Haumaitre et al., 2006). To generate ihPSCs, venous blood samples were obtained with informed consent from a family with inherited *HNF1B*-associated DKMs (Manchester Gene Identification Consortium Study (REC 11/H1003/3; IRAS ID 64321). Three siblings, each from a separate pregnancy, had bilateral DKMs detected on fetal ultrasonography. The female sibling had oligohydramnios and underwent termination because of a poor prognosis. Her two male siblings were born and were found to have ultrasound-bright kidneys, with loss of distinction between the cortex and medulla, core sonographic features of DKMs (Kohl et al., 2022). When assessed as young adults, their estimated glomerular filtration rates were modestly decreased (60-70 ml/min), and each had evidence of tubulopathy, e.g. hyperuricaemia, with one brother also having urinary glucose wasting. Each brother carries a deletion of exon 9 of *HNF1B* (del c.1654-? – c.1674-?; abbreviated as *HNF1B*^+/ΔExon9^) inherited from their father who has kidney disease and gout.

### hPSC cell culture

Stem cells were grown on culture plates coated with 5micro μg ml^-1^ recombinant human Vitronectin (rhVTN-N, Life Technologies, #A14700). HESCs were grown in mTeSR1 (StemCell Technologies, #85850) medium, whereas iPSCs were grown in TeSR-E8 (StemCell Technologies, #05990), with the medium changed every two days. The cells were passaged by treatment of the cultures with 0.5mM EDTA solution, pH8 (Invitrogen, #15575-038; diluted in PBS) and replating the cells in medium containing 5μM ROCK inhibitor, Y-27632 (Tocris, #1254) for 24h.

### CRIPSR/Cas9^n^ editing of hESCs

We first constructed a pair of plasmids expressing the nickase (D10A) version of Cas9 (Cas9^n^) and a gRNA each. The two gRNAs targeted the Cas9^n^ to nick at position 231 of the coding strand of *HNF1B* and at position 171 of the complementary strand, resulting in a deletion of 58 bases (plus/minus any indels) near the beginning of the coding sequence of the gene (Fig. 1A). Each gRNA-coding insert was designed as a pair of oligonucleotides that, upon annealing, produced overhangs that allowed cloning into the pX461 plasmid vector, (Addgene, #48140), under the control of a U6 promoter. The vector also expresses Cas9^n^, and a GFP tag for cell sorting. The primers used to create the inserts are listed in Table S2. Each pair of oligonucleotides was annealed and phosphorylated, using T4 Polynucleotide Kinase (NEB, #M0201S), then ligated into BbsI-digested pX461, to produce pX461-gRNA(HNF1B/231+) and pX461-gRNA(HNF1B/177-).

4×10^5^ MAN13 hESCs (Ye et al., 2017) were nucleofected with both pX461-gRNA(HNF1B/231+) and pX461-gRNA(HNF1B/177-), using the Amaxa™ P3 Primary Cell 4D-Nucleofector™ X Kit L (Lonza, #V4XP-3024), in 100μl nucleofection buffer according to the manufacturer’s instructions, on a Lonza 4D-Nucleofector (program DN100). 1ml of TeSR1 medium, containing 10 μM ROCK inhibitor, was then added to the cell mix and the cells were plated on a well of a 12-well plate coated with Vitronectin. The medium was replaced 16h later with 1ml fresh TeSR1 medium. Two days post-nucleofection, the cells were collected by TrypLE treatment (Life Technlogies, #12605-028) for 2min at 37°C and sorted for GFP fluorescence on a BD FACSAria Fusion flow cytometer. Two aliquots of the sorted, GFP+ cells of 5,000 and 10,000 cells were then plated in one Vitronectin-coated well each in a 6-well plate. The rest of the GFP+ cells (approximately 35,000) were plated on a separate well and – when confluent – they were stored in liquid N2, as backup. After 8-15 days, separate cell colonies had emerged on the sparsely seeded plates. Each was manually passaged, using a flame-pulled glass Pasteur pipette, into a separate well of a 24-well plate. Each clonal line thus produced was expanded further by EDTA passaging when confluent, and genomic DNA (gDNA) was extracted for genotyping using the Wizard® Genomic DNA Purification Kit (Promega, #A1120). A 781bp fragment around the expected mutation site was amplified by PCR, using Herculase II polymerase (Agilent, #600675) and was then sequenced to verify the presence/absence of the deletion. The primers used for PCR amplification and sequencing are detailed in Table S2.

### iPSC derivation

A total of 4 ml of blood were withdrawn and transferred to BD Vacutainer™ Hemogard closure plastic K2-EDTA tubes (BD 367525, BD). Tubes were inverted 10 times to ensure blood and EDTA were well mixed and kept at room temperature (RT) until processing. Peripheral blood mononuclear cells (PBMCs) were isolated by first mixing the each blood sample with an equal volume of PBS. An equal amount of Ficoll® Paque Plus (GE17-1440-02, Sigma) was then added in a 15 ml Falcon tube and the blood/PBS mix was carefully layered over the Ficoll reagent and centrifuged at 400g for 40 min, at RT. A total of 1ml PBMCs was isolated using a sterile Pasteur pipette and transferred into a new 15ml Falcon tube containing 9 ml of PBS. Diluted PBMCs were centrifuged at 200g for 10 min. The supernatant was discarded and the pellet was washed with 10 ml of PBS. Centrifugation was repeated at RT and the supernatant were discarded. PBMCs were resuspended in 2ml (0.5 ml medium per ml of starting blood) of Erythroid expansion medium, StemSpan™ SFEM II (#09605, STEMCELL Technologies) supplemented with StemSpan™ Erythroid Expansion Supplement (100X) (#02692, 50 STEMCELL Technologies). Typically, 1×10^6^ PBMCs were recovered per ml of blood. To expand the PBMC population, 5×10^5^ PBMC were plated in one well of a 6-well plate, containing 2 ml of erythroid expansion medium and incubated overnight. The following day, all non-adherent cells were transferred to a new plate and incubated overnight, while adherent cells were discarded. The cells were incubated and allowed to grow for a further 6 days, being fed every two days by carefully removing 1.5 ml of used erythroid expansion medium and replaced with fresh medium. The cells were then transduced with CytoTune™-iPS 2.0 Sendai Reprogramming Kit (A16517, Thermo Fisher), according to the manufacturer’s instructions. Briefly, per sample, 5×10^4^ cells were pelleted by centrifugation at 300g and were transduced by resuspension in 250micro μl erythroid expansion medium containing Sendai vectors (SeV) at a multiplicity of infection (MOI; viral particles per cell) of 5 for SeV-hKOS, 5 for SeV-hc-Myc and 3 for SeV-hKlf4. Cells were then centrifuged at 300g for 35 min to assist transduction, at RT, then resuspended and plated into two wells of a 24-well plate, where they were allowed to expand for a further four days, in a total volume of 600μl per well. The cells in each well were then transferred to a Vitronectin-coated well of a 6-well plate, containing 0.5 ml of ReproTeSR™ medium (#05921, STEMCELL Technologies). After two days, an extra 1ml of medium was added per well and the cells were allowed to settle down over the next 15 days, during which time iPSC colonies began to emerge. During this time, the medium was replaced with 1.5 ml ReproTeSR™ every 2 days. Colonies were manually cut into several pieces, using a pulled glass pipette, and each transferred to a Vitronectin-coated well of a 6-well plate. They were then passaged either manually, using a pulled glass pipette as above, or by EDTA treatment (see below), until approximately passage 20 before being used in experiments.

### 2D and organoid differentiation

The first differentiation steps occur in 2D over 7 days when dissociated cells are plated onto the air/medium interface of semipermeable membranes and maintained in 3D organoid culture for a further 18 days, unless otherwise stated. Differentiation of cells into kidney in 2D and 3D (organoids) was performed as described previously (Bantounas et al., 2018; Takasato et al., 2015), substituting APEL™ with APEL™2 medium, supplemented with 1% (v/v) PFHM-II Protein-Free Hybridoma Medium (Thermo Fisher Scientific, # 11370882), hereafter referred to as APEL™2. For 2D differentiation, stem cells were plated on Vitronectin-coated plates, at a density of 18,000 cells cm^-2^ in culture medium containing 10μM Y-27632. The following day the medium was replaced with STEMdiff™ APEL™2 (StemCell Technologies, #05270), containing 8 μM CHIR-99021 (Tocris, #4423) for three days, followed by APEL™2 supplemented with 200 ng ml^−1^ FGF9 (Peprotech, #100-23) and 1 μg ml^−1^ heparin (Sigma, #3149) for a further 10 days. Subsequently, the cells were cultured in basal APEL™2 medium which was changed daily. For 3D organoid cultures was performed according as described (Takasato et al., 2015). Cells were differentiated for the first 7 days as in the 2D protocol above, except that APEL/CHIR-99021 medium was used for the first four days. On day 7, cells were dissociated with TrypLE. Aliquots of 2×10^5^ cells were centrifuged at 400g for 2 min and the pellets were subsequently cultured on a medium-air interface by transfer onto a MilliCell cell culture insert (0.4μm pore size; Millipore, #PICM03050) inserted into the wells of 6-well plates. Upon transfer, the cells were exposed to APEL™2 medium containing 5μM CHIR-99021 for 1h, before reverting back to APEL™2 with FGF9/Heparin for a further 5 days. Finally, the 3D cultures were maintained in basal APEL™2 medium for up to another 13 days.

### RNA isolation from organoids and human embryonic tissue

RNA from kidney organoids was collected at day 0, 7, 12, 19 and 25 of the differentiation protocol. RNA was extracted using the miRVana miRNA isolation kit (Thermo Fisher, AM1560) according to the manufacturer’s instructions. Three organoids were pooled and lysed together to produce each sample. Four human felt kidneys (8, 8, 9 and 10 weeks of gestation) were obtained frozen and were homogenized on dry ice in Eppendorf tubes, using sterile plastic mini-pestles. Homogenized tissue was lysed in 600μl Lysis Buffer (provided with the miRVana kit) and RNA was then isolated using the miRVana kit according to the manufacturer’s instructions.

### Quantitative PCR

Quantitative real-time PCR was performed using the TaqMan® RNA-to-Ct™ 1-Step Kit (Thermo Fisher, #4392653) according to the manufacturer’s instructions, on a BioRad C1000™ Thermal Cycler fit with a CFX384™ Real Time System, using 15ng of RNA per reaction. The primers used are shown in Table S2.

### Next Generation RNA sequencing (RNAseq)

IBM13-08 and IBM13-19 hESCs were differentiated as described above in three independent experiments and RNA was collected as described above at days 0, 7 12, 19, 25 of differentiation. RNAseq was performed using an Illumina HiSeq4000 sequencer and unmapped paired-end sequences FastQC (http://www.bioinformatics.babraham.ac.uk/projects/fastqc/). Sequence adapters were removed, and reads were quality trimmed using Trimmomatic_0.39 (Bolger et al, 2014). The reads were mapped against the reference human genome (hg38) and counts per gene were calculated using annotation from GENCODE 36 (http://www.gencodegenes.org/) using STAR_2.7.7a (Dobin et al, 2013). Normalisation, Principal Components Analysis, and differential expression was calculated with DESeq2_1.28.1 (Love et al, 2014).

### GO and KEGG pathway analysis

Gene ontology enrichment was studied using Enrichr v3.1 (PMID: 27141961). Enriched KEGG and Reactome pathways were identified using the ClusterProfiler package (Yu et al., 2012) and visualized using the Pathview package (Luo and Brouwer, 2013). For Gene Set Enrichment Analysis (GSEA), a pre-ordered list of log2 fold changeFC values, and for Over Representation Analysis (ORA), an input list of DEGs was used to get enriched pathways. Up-regulated pathways were defined by a normalized enrichment score (NES) > 0 and the down-regulated pathways were defined by an NES < 0. Pathways with BH adjusted-P value ≤ 0.05 were chosen as significantly enriched.

### In silico identification HNF1B-regulated promoters

Transcription factor (TF), HNF1B-responsive genes were searched using iRegulon (Janky et al., 2014). iRegulon detects the TFs and their targets by scanning known TF-binding promoter motifs as well as the predicted motifs discovered from the Encyclopedia of DNA Elements (ENCODE) Project chromatin immunoprecipitation-sequencing data. It includes 1,121 human regulatory tracks with ChIP-seq data for 247 sequence-specific TFs across 43 different cell types and conditions. We selected 20-kb upstream parameter for the options "Putative regulatory region," "Motif rankings database," and "Track rankings database" to identify the targets. Motif sequences of HNF1B were explored with in silico predicted and experimentally validated DNA-binding motifs collection databases (JASPAR2018_CORE_vertebrates_non-redundant, Jolma 2013, TRANSFAC, UniPROBE mouse, HOCOMOCO v11 and Swiss Regulon) using TOMTOM (Gupta et al., 2007) (e-value 0.5; min overlap between motifs).

### Western blotting

Per sample, three organoids were pooled and homogenised in 350 µL of RIPA lysis buffer (Thermo fisher Scientific, #89900) with protease and phosphatase inhibitors. Twenty to thirty μg of sample was loaded per well, on a NuPAGE^TM^ 10% Bis-Tris, 1.0 mm, Mini protein Gels (10-well) (Thermo fisher Scientific, #NP0301) and electrophoresed in 1x MES buffer (diluted from a 20X stock; Thermo fisher Scientific, #NP0002). Proteins were transferred onto a nitrocellulose membrane in the iBlot2 transfer stack (Thermo fisher Scientific, #IB23001) using an iBlot 2 Dry Blotting System (Life Technologies). Membranes were blocked with 5% skimmed milk powder diluted in PBS-Tween 0.1% (v/v) (Sigma-Aldrich, #PP9416) (PBS-T), for 1 hour at room temperature with agitation. Primary antibodies (Table S3) were diluted in blocking solution and incubated with the membrane, at 4°C with agitation overnight. The membrane was then washed in PBS-T 3 times for 10 minutes and secondary antibodies (Table S3), diluted in blocking solution, were incubated for 1 hour at room temperature, followed by three final 10-minute PBS-T washes. Membranes were imaged with the Odyssey CLx Imaging system (LI-COR Biosciences, Germany) and densitometry was performed with the Image Studio Lite quantification software.

### Immunocytochemistry of 2D cultures

Cells were washed twice in PBS and then fixed in 4% paraformaldehyde (PFA) for 20 minutes, followed by another two PBS washes. The fixed cells were blocked and permeabilised for 30 min with 3% bovine serum albumin (BSA)/0.3% Triton-X in PBS before overnight incubation at 4°C with primary antibodies (Table S3) diluted in 3% BSA/PBS. They were then washed three times with PBS/0.1% Triton-X, followed by Alexa-Fluor™-488- or Alexa-Fluor™-594-labelled, species-specific secondary antibodies (Life Technologies; 1:300 dilution in 3% BSA/PBS). Images were collected on a Zeiss Axioimager.D2 upright microscope using a 63x/Plan-neofluar objective and captured using a Coolsnap HQ2 camera (Photometrics) through Micromanager software v1.4.23. Images were then processed and analysed using Fiji-ImageJ (http://imagej.net/Fiji/Downloads).

### Immunohistochemistry of organoid sections

Organoids were fixed in 4% paraformaldehyde, embedded in paraffin and sectioned at 5 µm. Sections were dewaxed and rehydrated, and alternate slides were stained with haematoxylin and eosin (H&E) to assess overall tissue architecture. Images were acquired on a 3D-Histech Pannoramic-250 microscope slide-scanner using a x20 objective (Zeiss) and selected images were captured using the Case Viewer software (3D-Histech). After rehydration, other slides were boiled in an 800W microwave in 10 mM sodium citrate buffer (pH 6.0). After cooling to room temperature, endogenous peroxidase activity was blocked using 0.3% H2O2 in PBS for 10 minutes. Sections were permeabilized using 0.2% Triton X-100 (Sigma-Aldrich) for 10 minutes and blocked using 1% bovine serum albumin (BSA) with 10% serum from the species in which the secondary antibody was raised. Sections were incubated overnight at 4°C with the primary antibody + 1% BSA. Primary antibodies used are listed in Table S3. Biotin-conjugated species specific secondary antibodies with 1% BSA were incubated at room temperature for 2 hours. Following PBS washes, slides were incubated in avidin-biotin enzyme complex (Vector Laboratories VECTASTAIN Elite ABC Reagent, PK-6100) for 1 hour at room temperature. Peroxidase activity was detected with the 3, 3’-diaminobenzidine (DAB) peroxidase substrate solution (Vector Laboratories, SK4100) in some cases with haematoxylin counterstain. Sections were dehydrated and mounted with DPX mounting medium and examined under a Leica DMLB 2 microscope. Negative controls omitted primary antibodies. For immunofluorescent imaging, anti-rabbit Alexa-Fluor™ 488 (Thermo Fisher Scientific, # A11034) and anti-mouse Alexa-Fluor™ 594 (Thermo Fisher Scientific, #A11032) secondary antibodies and DAPI nuclear stain were used. Sections were mounted with Vectashield antifade mounting medium (Vector Laboratories, #H-1000). Images were acquired using an Olympus BX63 upright microscope using a DP80 camera (Olympus) through CellSens Dimension v1.16 software (Olympus) and slide scanned on a 3D-Histech Pannoramic-250 microscope slide-scanner using a 40x/0.95 Plan Apochromat objective (Zeiss) and captured using the Case Viewer Pannoramic250 slide scanner software (3D-Histech) at the University of Manchester Bioimaging facility, and processed and analysed using Fiji-ImageJ (http://imagej.net/Fiji/Downloads).

### Immunohistochemistry and periodic acid-Schiff (PAS) staining of fetal kidneys

Sections were dewaxed and rehydrated, endogenous peroxidase activity was blocked using 0.3% H2O2 in PBS for 20 minutes with agitation. Sections were then boiled in a microwave in 10micro mM sodium citrate buffer (pH 6.0). After cooling to room temperature, they were incubated with saturated lithium carbonate cat. 26684-03 (Generon Ltd) in distilled water for 30 minutes, permeabilised with Tris-buffered saline (TBS)/triton X-100 0,3% for 20 minutes followed by incubation in blocking buffer for 45 minutes in a humidified chamber at room temperature. Primary antibody GRIK3 (table S3) was diluted in 1:800 and tissues were incubated overnight at 4°C. After washing, biotin-conjugated secondary antibody was incubated at room temperature for 1 hour. Following washes, slides were incubated in avidin-biotin enzyme complex (above) for 30 minutes at room temperature and peroxidase activity was detected as above. Periodic acid-Schiff stain (PAS) was carried out according to manufacture instructions using the kit 395B (Scientific Laboratory Supplies Ltd) as previously described (Yuan at al., 1999). Images were captured as above.

### *In situ* hybridisation

Organoids were fixed in 4% paraformaldehyde and sectioned at 5 μm. BaseScope (ACDBio, Newark, CA, USA) was adapted from Lopes et al 2009 and following the manufacturer’s instructions using the BaseScope detection reagent Kit v2-RED. RNA (red) was detected using Fast RED and nuclei counterstained with Gill’s haematoxylin. The following custom-made BaseScope probes were used: BA-Hs-*HNF1B*-3zz-st targeting 1078-1252; BA-Hs-*HNF1A*-3zz-st targeting 1453-1565 and BA-Hs-*GRIK3*-No-XMm-3zz-st, targeting 3258-3414 and the following control probes: human (HS)-*PPIB-3zz* cat. 701031 (positive control); and *DapB* (bacterial gene)*-3ZZ* cat.701011 (negative control). The positive control probe was the widely expressed *PPIB* transcript encoding peptidylprolyl isomerase B, and a negative control probe was *DapB* encoding 4-hydroxy-tetrahydrodipicolinate reductase from the *Bacillus subtilis* soil bacterium, a gene that is absent in mammals. Images were acquired using an Olympus BX63 upright microscope using a DP80 camera (Olympus) through CellSens Dimension v1.16 software (Olympus).

### cAMP-induced tubule dilatation

We used a previously described protocol used for intact metanephroi maintained in organ culture (Anders et al., 2013). The growth medium of organoids was supplemented with 100μM with 8-Bromoadenosine 3’,5’-cyclic monophosphate sodium salt (8-Br-cAMP) (Sigma, #B7880) in iPSC organoids or 200μM forskolin (FSK) (Tocris, #1099) in hESC organoids, starting on day 14 of the protocol. Treatment continued until day 25 (iPSC organoids) or day 32 (hESC organoids). Dilatations were identified by their translucent/bright appearance under phase microscopy. Their number and size were measured using Fiji-ImageJ.

### Cell Proliferation Assay

Day 25 differentiated kidney organoids were incubated with 10 µM of 5-Bromo-2’-deoxyuridine (BrdU) (Sigma-Aldrich, #B5002-100MG) in basal medium for 2 hours at 37°C, by placing 1.2ml BrdU supplemented medium underneath the transwell filter and 1ml inside the transwell. Organoids were then fixed and processed and immunohistochemistry was used to stain BrdU positive nuclei.

## ACKNOWLEDGEMENTS

We acknowledge research support from: Kidney Research UK project grant JFS/RP/008/20160916 (SJK, ASW and IB); Medical Research Council Project grant MR/T016809/1 (ASW and FML); Engineering and Physical Sciences Research Council (EPSRC)/Medical Research Council (MRC) Centre of Doctoral Training grant EP/L014904/1 (KR); European Union (SYBIL European Community’s Seventh Framework Program, FP7/2007-2013, 602300 (SJK, SW), Wellcome Leap Human Organs Physiology and Engineering (HOPE) Initiative (ASW, SJK and IB); Kidneys for Life pump priming projects 2021 (ASW, SJK, IB and KMR). Support is also acknowledged from: the Malaysian Ministry of Higher Education and International Islamic University Malaysia (FT) and an Erasmus Scholarship (SH).

## AUTHOR CONTRIBUTIONS

SJK, ASW, IB and KMR designed the studies, supervised the project and drafted the paper. IB, KMR, FT, FML, SW, NB, LW and SH undertook laboratory experiments. SC provided kidneys for histology. KAH and ASW assessed the *HNF1B* family and sourced samples to generate hiPSCs. LAHZ and SYK undertook bioinformatic analyses. All authors discussed the results and commented on the manuscript.

## SUPPLEMENTAL FIGURE LEGENDS

**Supplemental Figure 1.**
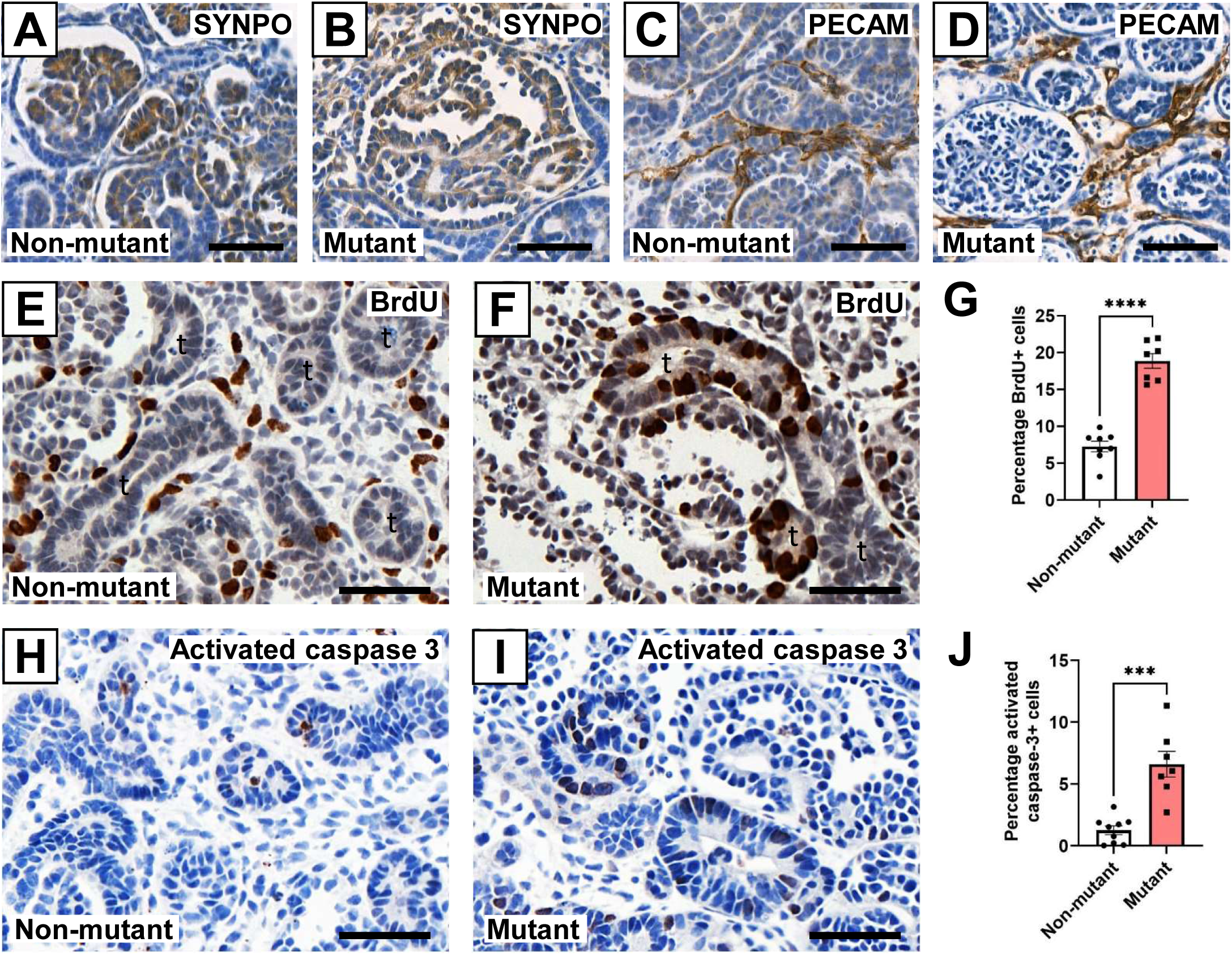
Glomerular histology and cell turnover in hESC-derived organoids. Synaptopodin (SYNPO) immunostaining (brown) of glomeruli in non-mutant (**A**) and *HNF1B* heterozygous mutant (**B**) organoids with haematoxylin counterstain (blue). Mutant glomeruli contained less compact podocyte tufts. PECAM1 immunostaining (brown) in non-mutant (**C**) and mutant (**D**) organoids with haematoxylin counterstain (blue) showed capillary-like structures between tubules. BrdU immunostaining (brown) marking proliferative cells in non-mutant (**E**) and mutant (**F**) organoids with haematoxylin counterstain (blue). Tubules indicated by *t*. **(G)** Quantification showed a significantly increased percentage of BrdU+ nuclei in mutant compared with non-mutant tubules (mean±SEM; n=8 non-mutant and n=7 mutant organoids across three independent differentiation experiments; ****p<0.00005, t-test). Activated caspase 3 immunostaining (brown) marking cells undergoing apoptosis in non-mutant (**H**) and mutant (**I**) organoids with haematoxylin counterstain (blue). **(J)** Significantly increased percentage of activated caspase 3 immunostained cells in mutant compared with non-mutant organoids (mean±SEM; n=9 non-mutant and n=7 mutant organoids across three independent differentiation experiments; ***p<0.0005, t-test). Bars are 50 μM.

**Supplemental Figure S2.**
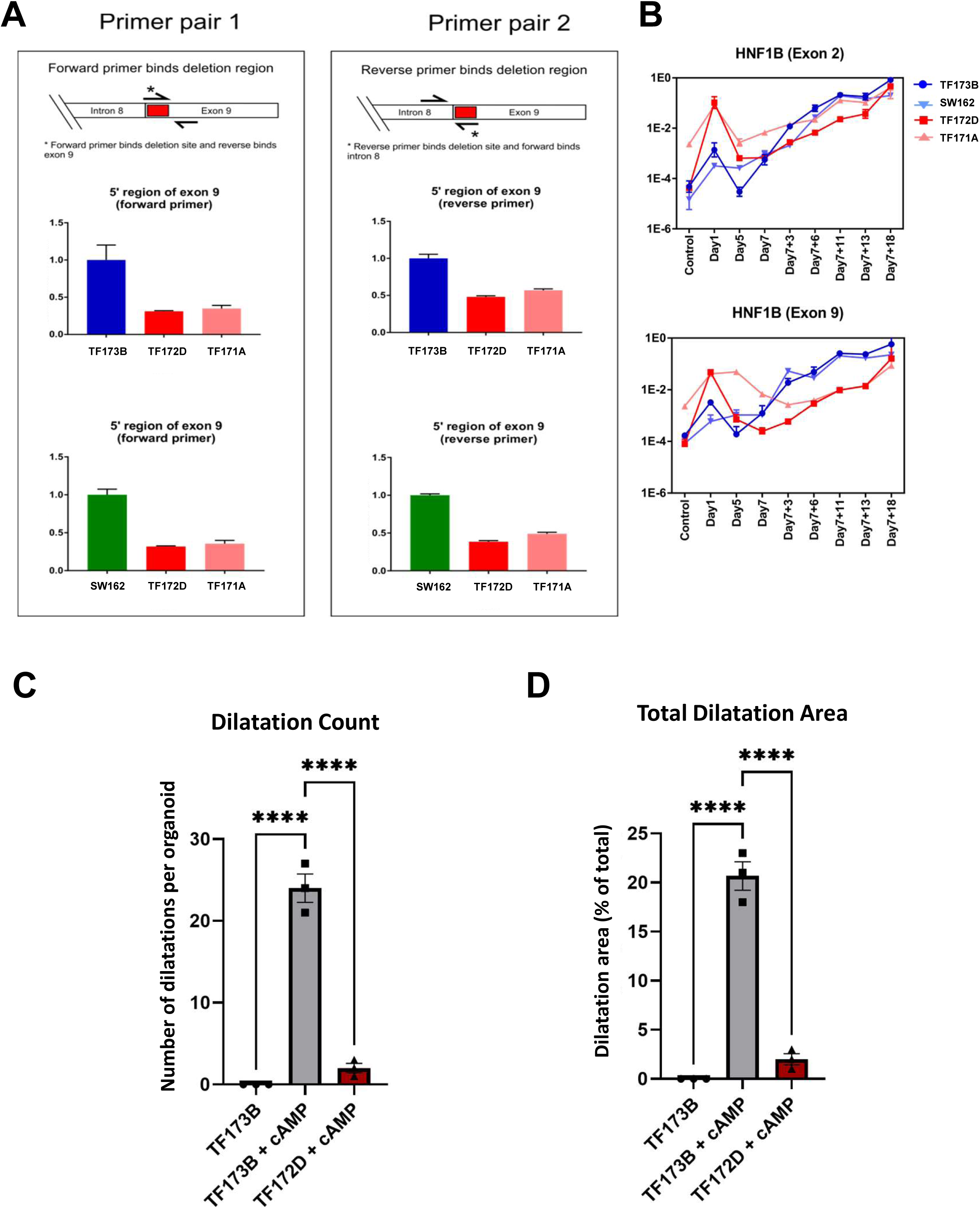
Characterisation of patient-derived iPSC lines. **(A)** qPCR of genomic DNA from *HNF1B^+/ΔExon9^*iPSCs (TF171A and TF172D) and unaffected control *HNF1B^+/+^* iPSCs (TF173B and SW160) with two primer pairs binding to the predicted deletion site in exon 9. A halving of gene dosage in TF171A and TF172D was detected compared with the *HNF1B^+/+^* mother (TF173B) and an unrelated control (SW160), confirming the heterozygous nature of the mutation (mean±SEM from 3 technical replicates). **(B)** qPCR time course of *HNF1B* mRNA levels, detected using either exon 2 (top) or exon 9-specific (bottom) primers. The former would detect *HNF1B* mRNA generated by wild-type and mutant alleles, whereas the latter detects only wild-type mRNA. Two *HNF1B^+/+^* iPSC lines were used for comparison, TF173B (mother) and SW160 (an unrelated control). Transcript levels detected by exon 9 primers were lower in mutants than in controls during the organoid phase (*day 7+3* onwards). **(C)** Quantification of numbers of dilated structures per organoid in *HNF1B^+/+^* (TF173B) and *HNF1B^+/ΔExon9^* (TF172D) organoids following 8-Br-cAMP exposure showing markedly lower numbers of dilated structures in *HNF1B^+/ΔExon9^*organoids (n=3 organoids across three differentiation experiments; p****<0.00005, one-way ANOVA with multiple comparisons). **(D)** Quantification of percentage dilatation area per organoid in *HNF1B^+/+^* (TF173B) and *HNF1B^+/ΔExon9^* (TF172D) organoids following 8-Br-cAMP exposure. Note significantly decreased percentage area of dilated structures in *HNF1B^+/ΔExon9^* organoids (n=3 organoids across 3 independent differentiation experiments: p****<0.00005, one-way ANOVA with multiple comparisons).

**Supplemental Figure S3.**
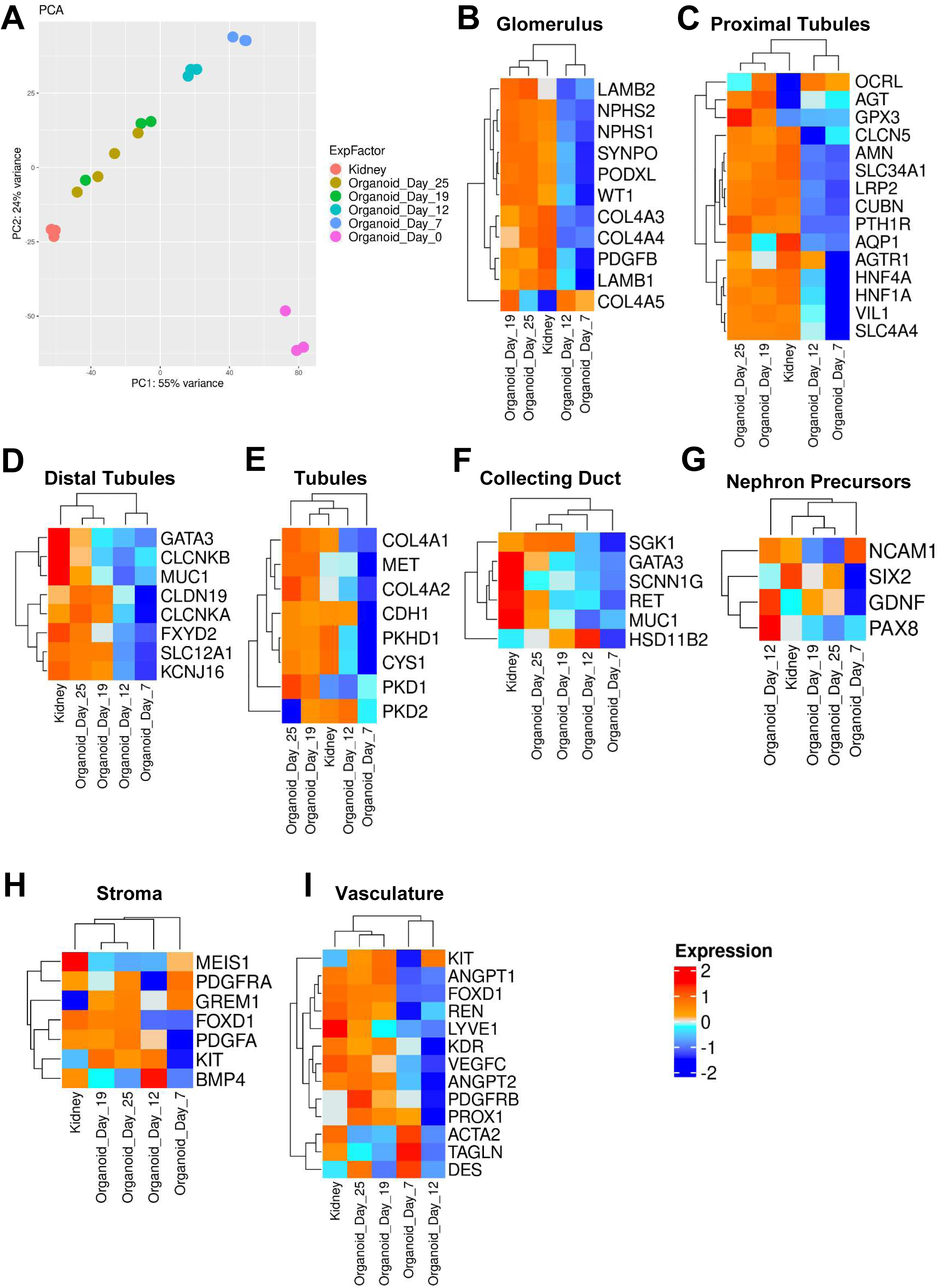
Transcriptomic profiling of differentiating non-mutant kidney organoids in comparison to fetal kidney. **(A)** PCA of RNAseq of differentiating wild-type hESCs. As differentiation proceeded from PSCs (*Organoid day 0*), profiles approached those of 8-10 week human fetal kidneys. Profiles of day 19 and 25 organoids were similar, suggesting that this period represents a plateau of *in vitro* differentiation when the whole transcriptome is considered. **(C-G)** Heat maps of gene expression marking specific cellular kidney compartments. Glomerular and PT transcripts rose during organoid maturation so that by days 19 and 25 they resembled those in fetal kidneys. These genes include *CUBN* and *LRP2*, whose protein products Cubilin and Megalin, were studied in organoids in Fig. 3. While some DT and CD genes were expressed, they tended to remain lower than in the fetal kidney. The *Tubule* panel showed several kidney epithelial genes whose expression was not limited to particular nephron segments or CDs. Several *Nephron precursor* genes are expressed in organoids and tended to be downregulated by day 25. Organoids also expressed several genes characteristic of blood and lymphatic vessels (*Vasculature*) and stromal cells. Each point of the heat map represents the mean of three (days 7-19) or four (‘Kidney’ and day-25) independent differentiation experiments.

**Supplemental Figure S4.**
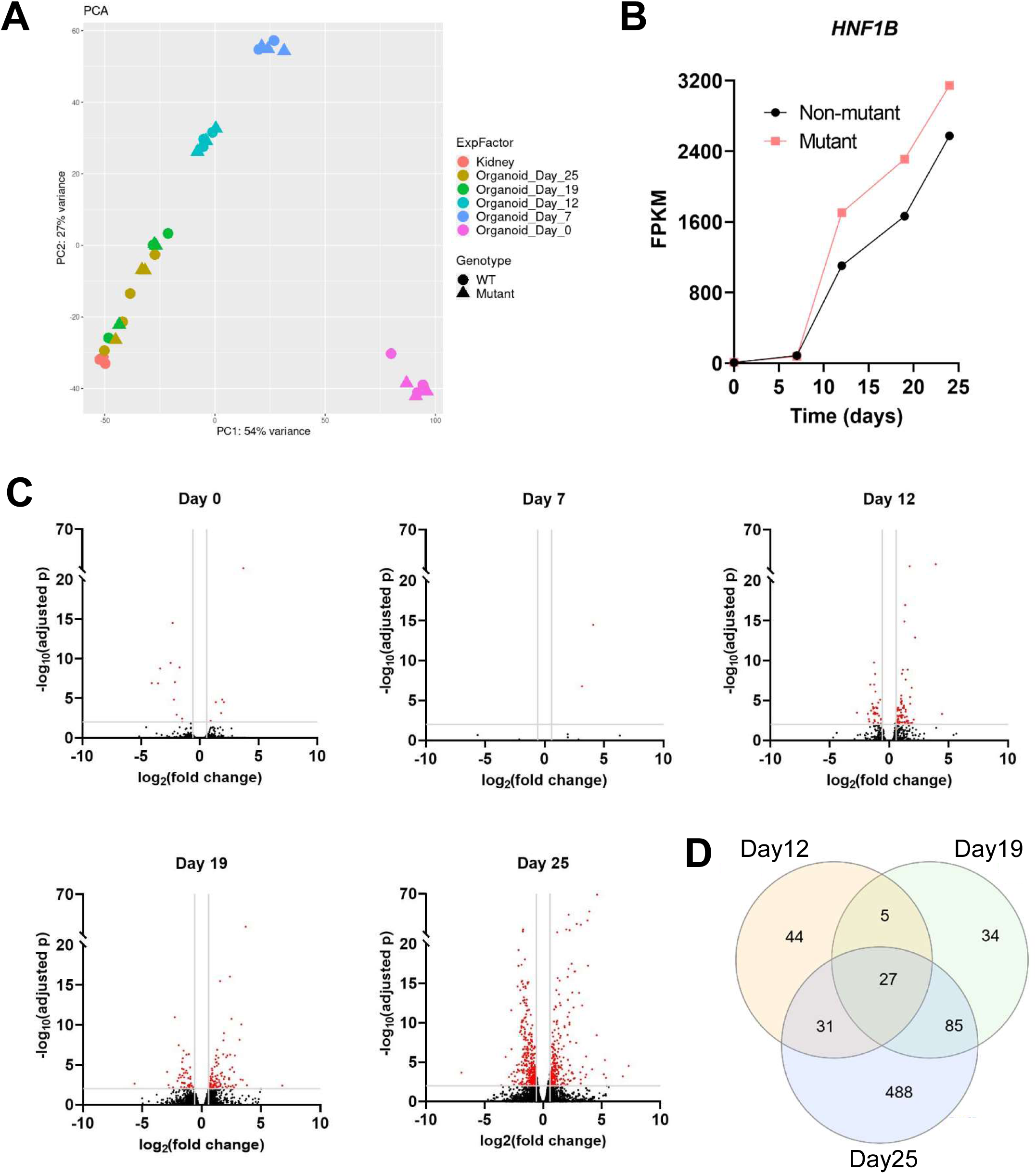
Transcriptional profiles of heterozygous *HNF1B* mutant cells compared with isogenic wild-type controls during differentiation to organoids. (A) PCA showing that progression with over the 25 day protocol was similar in the two genotypes. (B) *HNF1B* transcripts in the bulk RNAseq increased similarly in each genotype. (C) Volcano plots demonstrating the progressive increase in numbers of significantly (red) differentially expressed genes during the organoid phase (*Day 12* onwards). (D) Venn diagram of significantly differentially expressed transcripts in the organoid phase of culture.

**Supplemental Figure S5.**
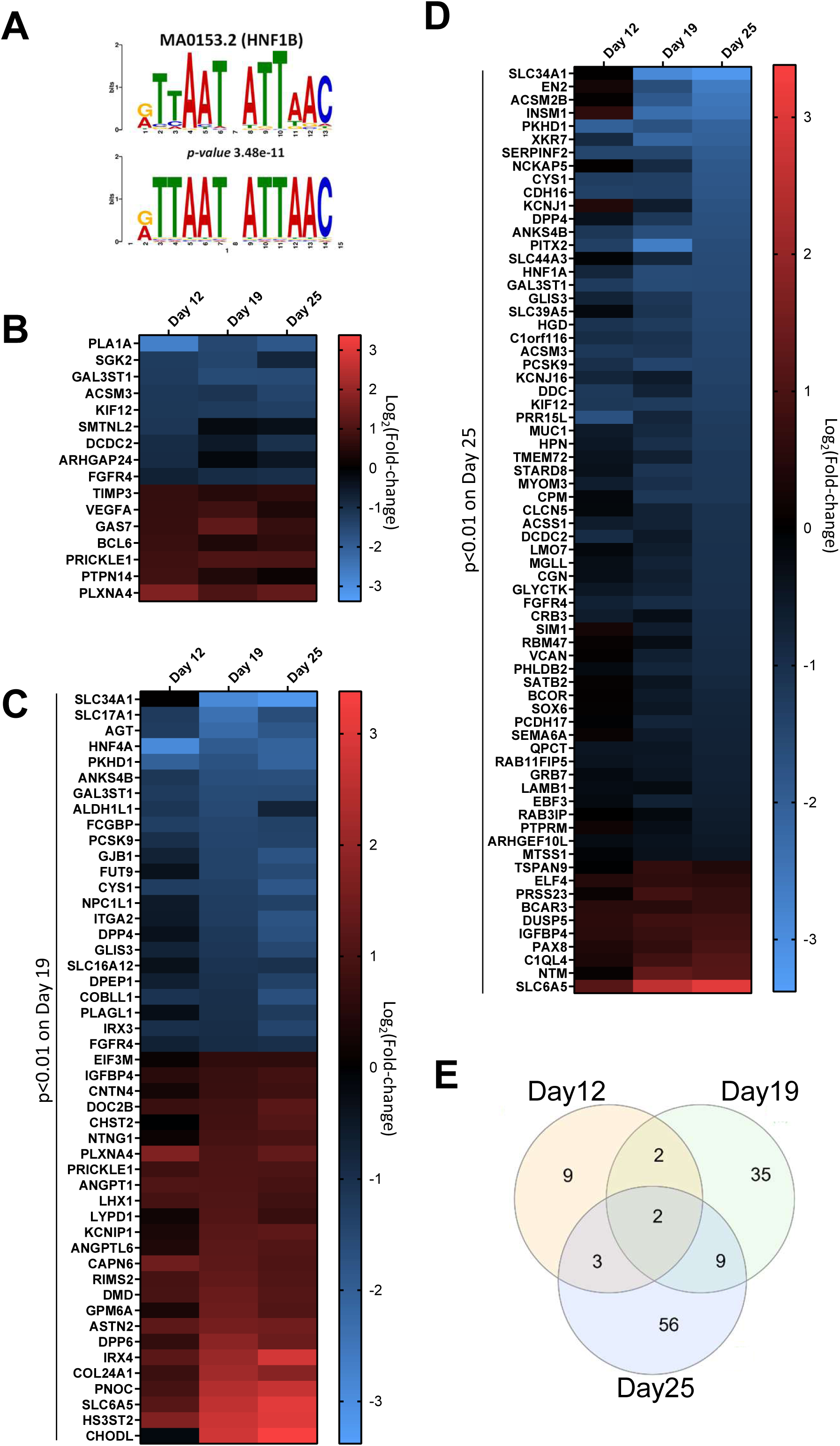
Differentially expressed genes containing the canonical HNF1B binding site. **(A)** Top: Sequence motif logo of HNF1B (MA0153.2; p-value 3.48× 10^−11^) transcription factor, within significant differentially expressed genes (padj≤0.05), created by TOMTOM from JASPAR2018_CORE_vertebrates_non-redundant database. Bottom: canonical *HNF1B* binding site used in this analysis.. **(B-D)** Heatmaps showing the difference between non-mutant and *HNF1B* mutant organoids, in expression levels of genes that contain the *HNF1B*-binding consensus sequence in their promoters, on day (B) 12, (C) 19 and (D) 25 of differentiation. Note that on days 12 and 19 an approximate equal number of genes were down-(blue) or up- (red) regulated, whereas on day 25 the majority of differentially expressed genes were downregulated. **(E)** Venn diagram quantifying the numbers of significantly differentially expressed genes at three stages of organoid culture.

**Supplemental Figure S6.**
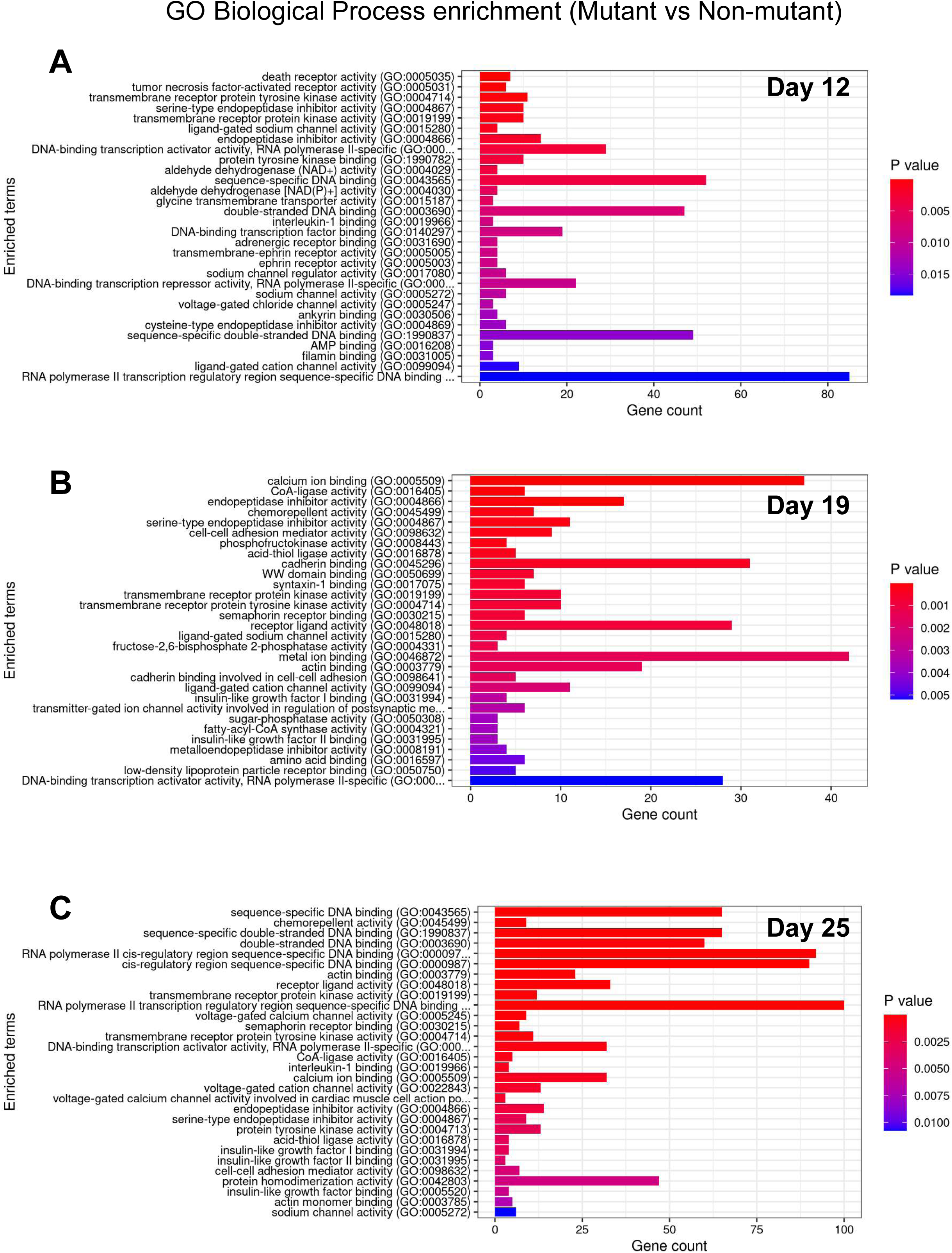
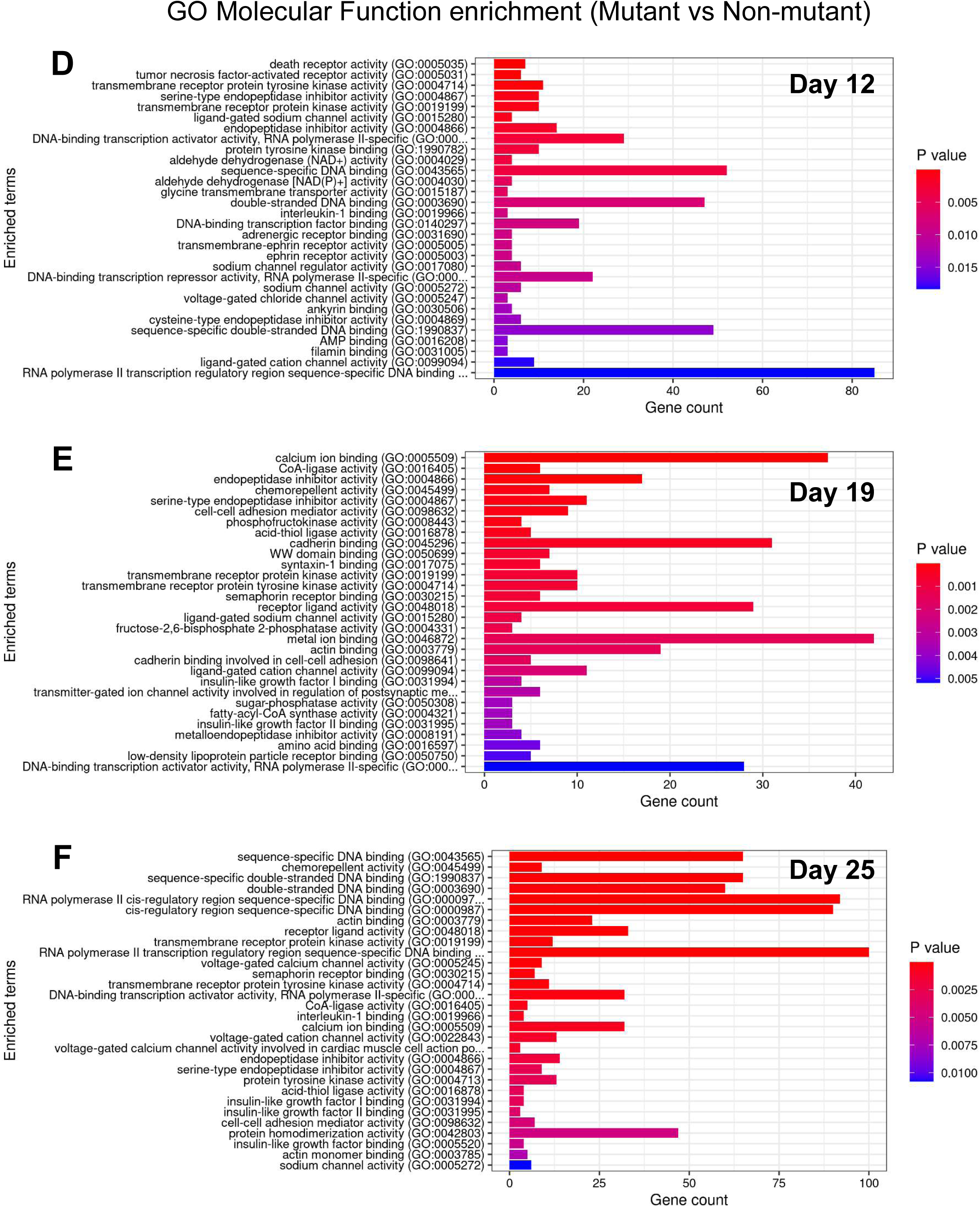
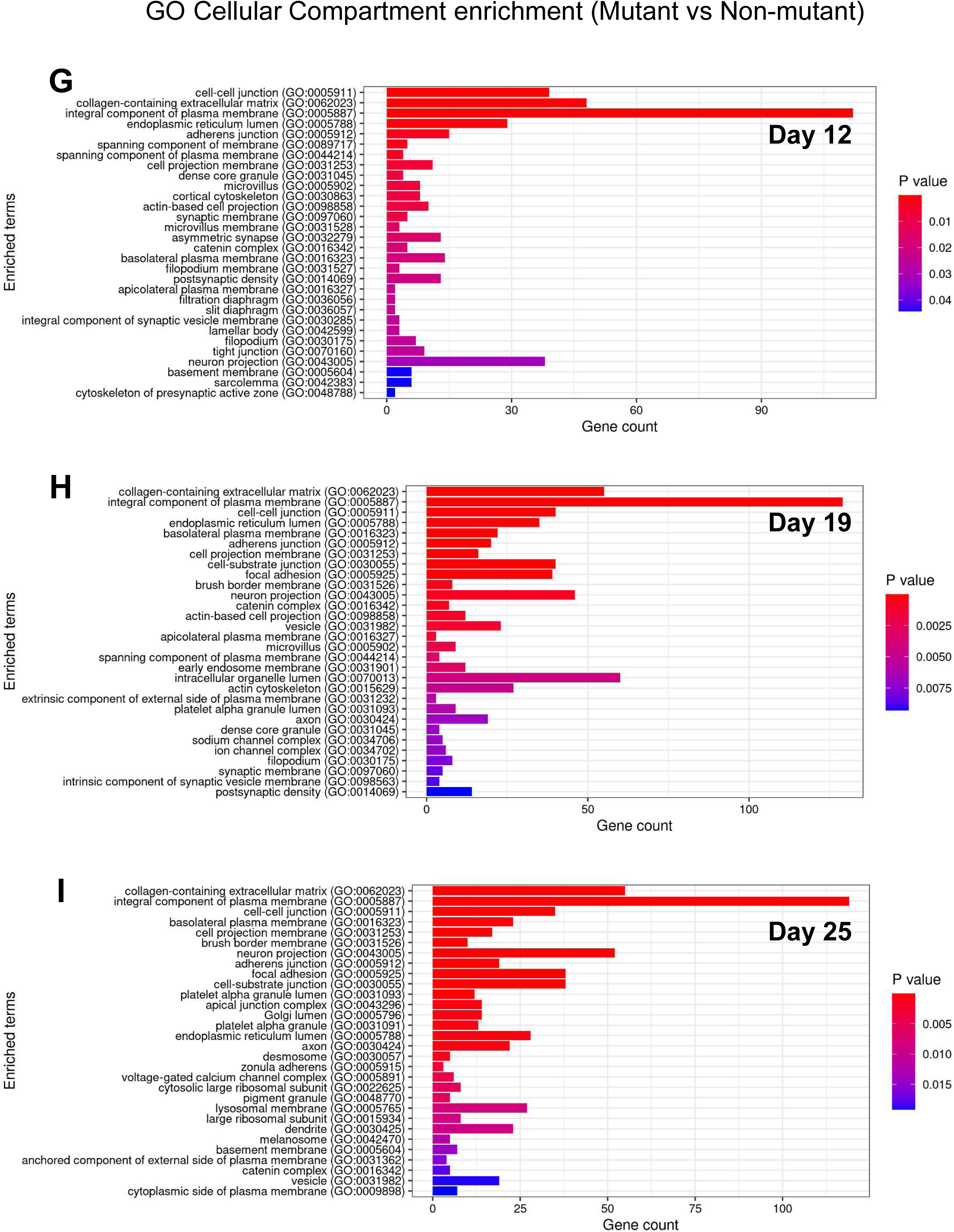
GO terms enrichment analysis comparing isogenic non-mutant and *HNF1B* mutant organoids across organoid development. **(A-I)** Bar charts of terms enriched in mutant organoids compared with non-mutant on days 12, 19 and 25 of differentiation. Graphs depict *biological process* (A-C), *molecular function* (D-F) and *cellular compartment* (G-I). Red bars indicate upregulated in mutant organoids, and blue indicate downregulated in mutant organoids.

**Supplemental Figure S7.**
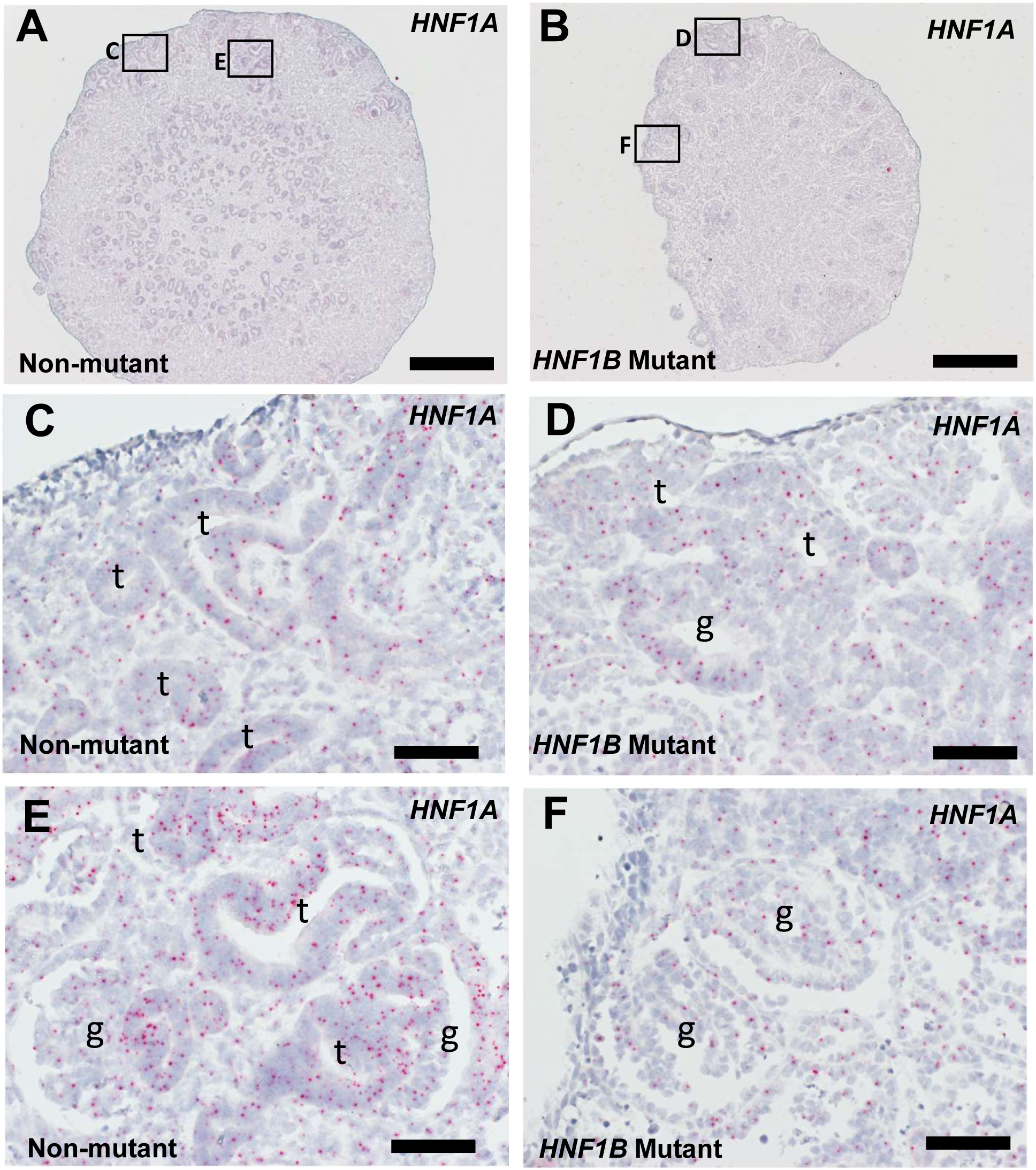
H*N*F1A transcript localisation in organoids. *HNF1A* Basescope ISH was performed on histological sections of kidney organoids at day 25. Signals appear as red dots; nuclei were counterstained blue with haematoxylin. **(A-B)** Low power overview of non-mutant and *HNF1B* mutant organoid. **(C-F)** High power frames of boxes indicated in (A-B). Note generally decreased *HNF1A* expression in mutant tubules (D) compared with non-mutant tubules (C, E). “t”, tubules; “g”, glomeruli. Scale bars: (A-B) 200 μM; (C-F) 20 μM.

**Supplemental Figure S8.**
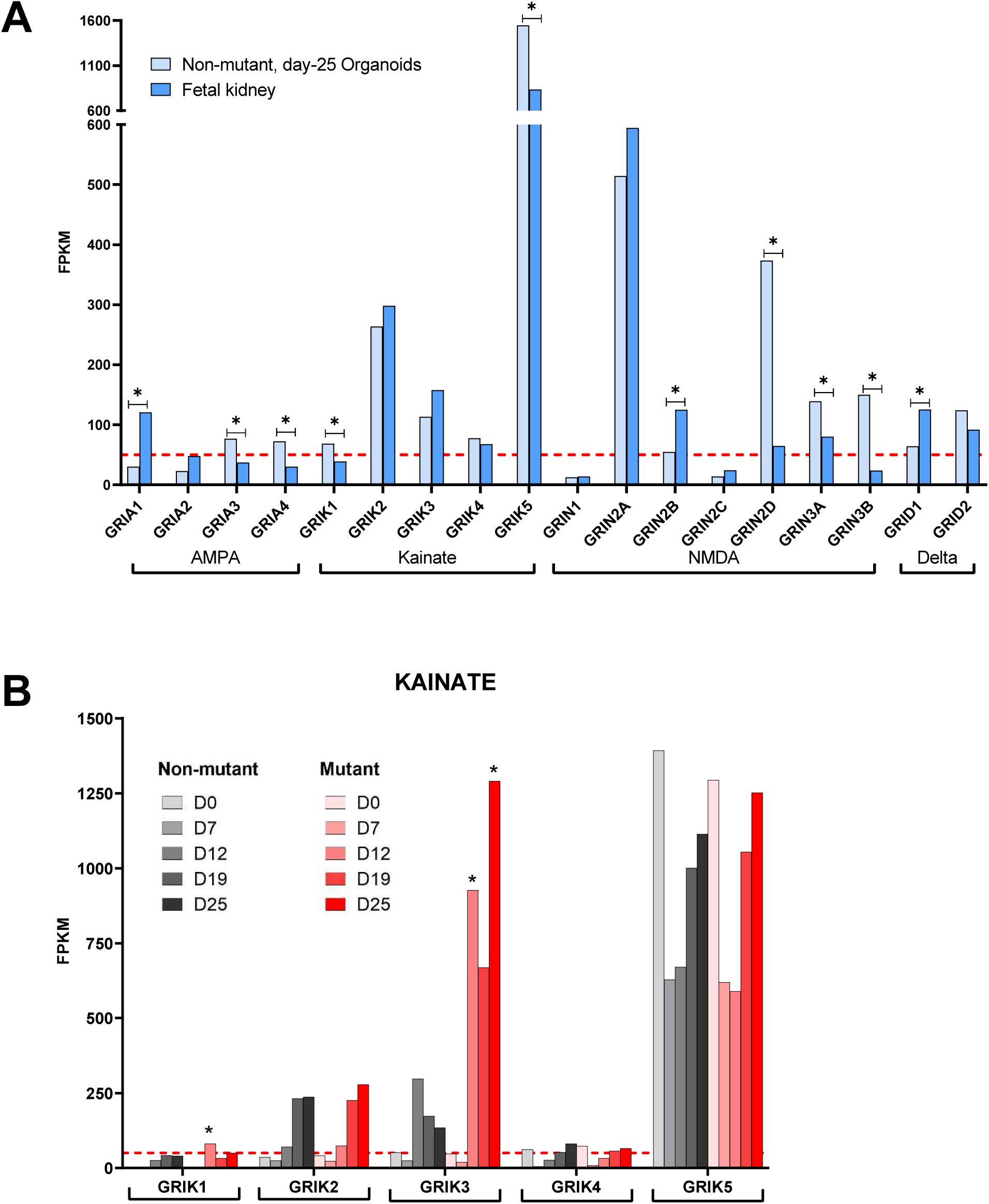
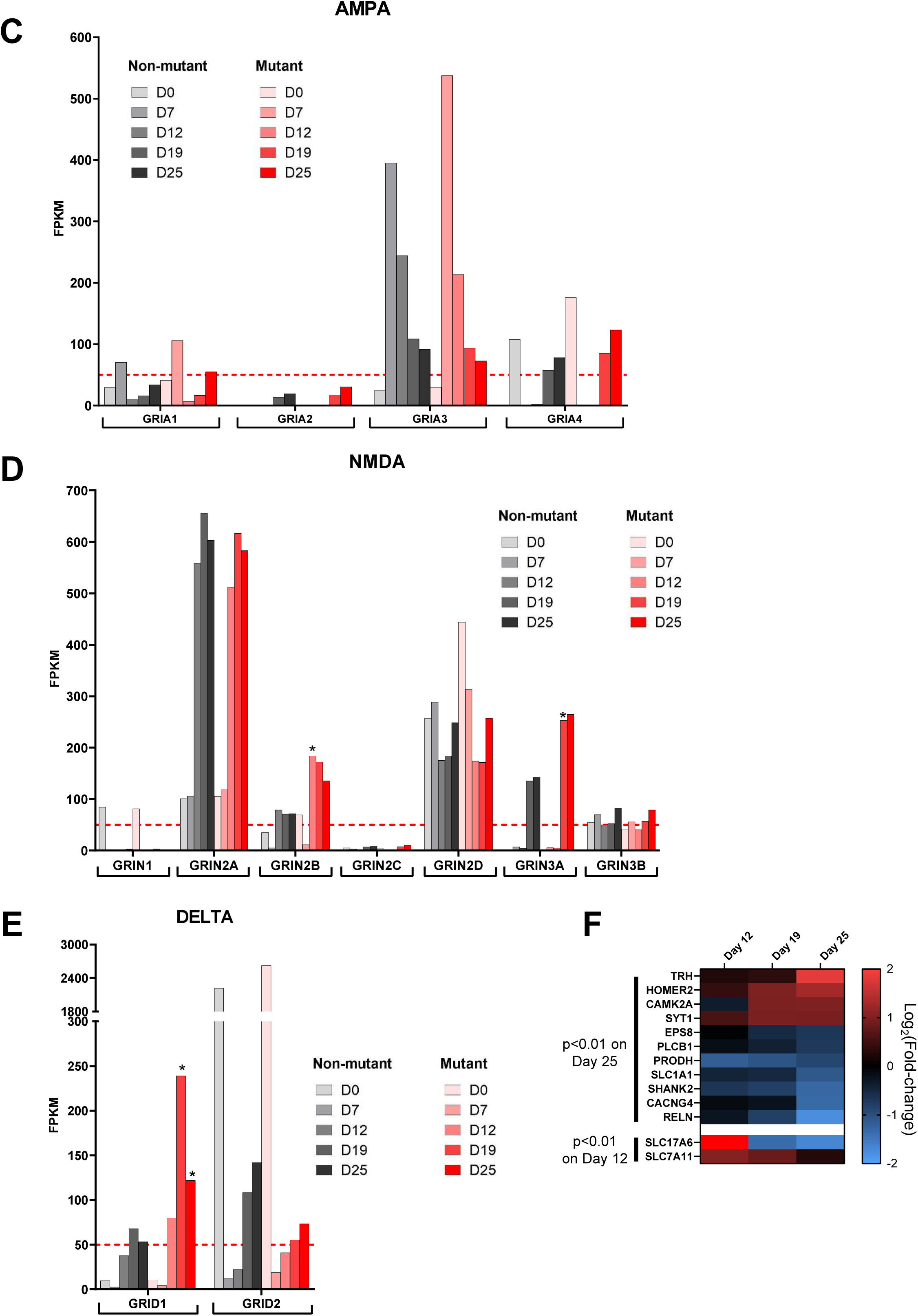
Expression of glutamate receptors (GluR) in human fetal kidney and hESC-derived kidney organoids. **(A)** GluR subunit expression (average reads from bulk RNAseq) in non-mutant day 25 organoids (light blue) and human fetal kidneys (dark blue). In general, expression levels were of similar magnitude between organoids and native kidneys, with some differences indicated (n=4 independent differentiation experiments, *adjusted p<0.05). **(B-E)** Time course of expression of members of different families of GluR subunits in non-mutant (*grey/black)* and *HNF1B* mutant (*pink/red*) organoids. Differences between time-matched mutant and non-mutant organoids are indicated (n=3 independent differentiation experiments, *adjusted p<0.05). **(F)** Heatmap of genes belonging to the GO pathways term Glutamate Signalling (excluding GluR themselves), with significantly different (adjusted p<0.01) expression in *HNF1B* mutant organoids, in our RNAseq dataset.

**Supplemental Figure S9.**
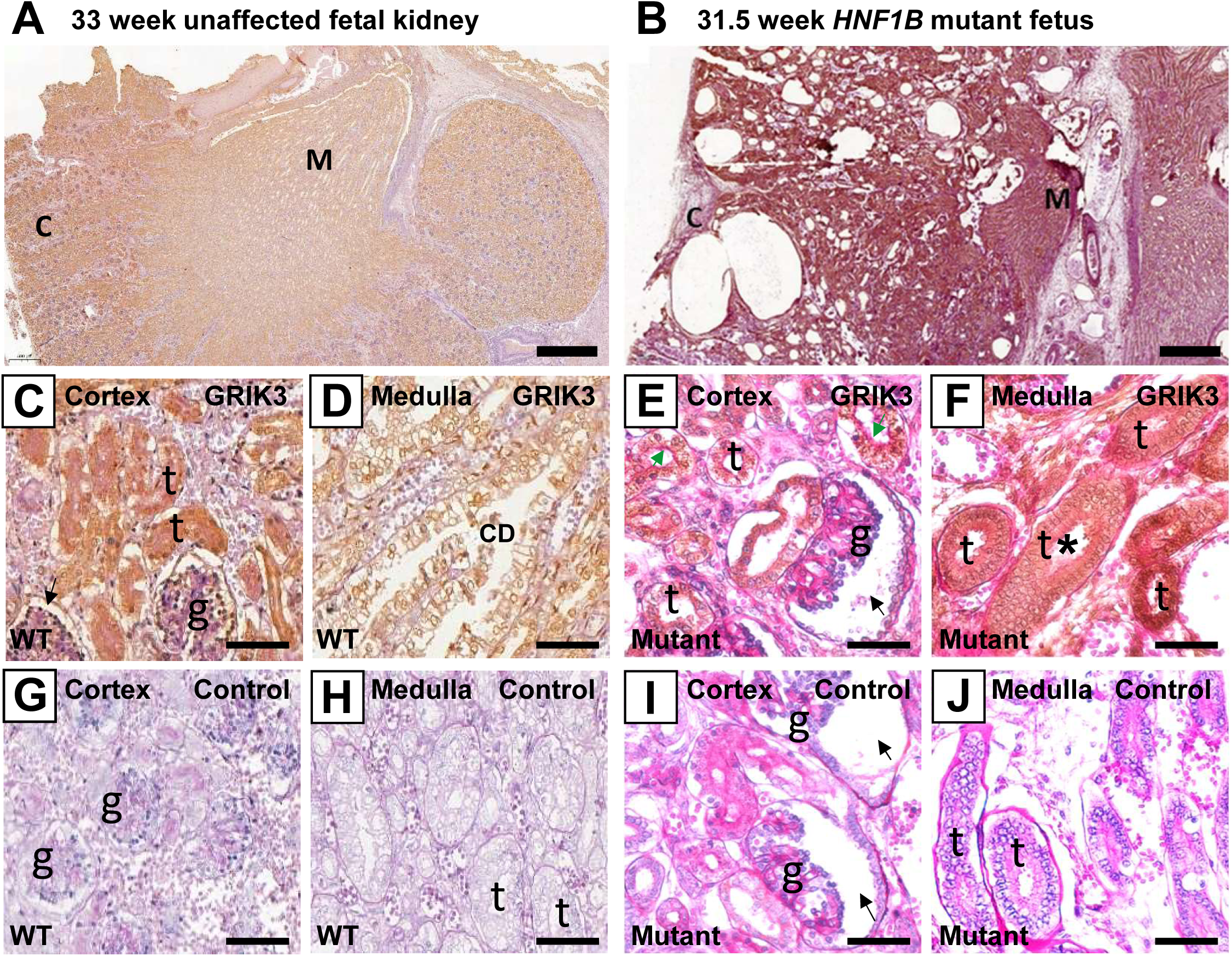
GRIK3 immunostaining of human fetal kidneys. A, C, D, G and H are sections of a control third trimester kidney, and B, E, F, I and J are from a third trimester DKM from a fetus with a heterozygous frameshift in exon 7 of *HNF1B*. All panels were counterstained with haematoxylin (blue nuclei) and PAS (pink basement membrane, tubule brush border and interstitial scarring). A-F were also immunostained for GRIK3 (brown) and in G-J the primary antibody was omitted. **(A)** Low power overview of the control kidney; note the normal arrangement of the cortex (C) and medulla (M). **(B)** Low power overview of the *HNF1B* mutant kidney; note the disorganised arrangement of cortex (C) and medulla (M) and cysts. **(C)** GRIK3 immunostaining in cortical tubules (t) and Bowman capsule (arrowhead) in the control kidney. **(D)** GRIK3 immunostaining in a branched collecting duct (CD) in the control kidney. **(E)** Mutant kidney contained glomeruli with dilated Bowman spaces (black arrow) and GRIK3 immunostaining in large cortical tubules (t) that contained PAS+ brush border (green arrows), suggesting they are PT-like tubules. **(F)** Other areas of the mutant DKM were rich in multi-layered dysplastic tubules (t with asterisk in lumen) and that immunostained for GRIK3. **(G-J)** are similar views to C-F but with primary antibody omitted. Scale bars: (A-B) 500 μM (C-J) 40 μM.

**Supplemental Figure S10.**
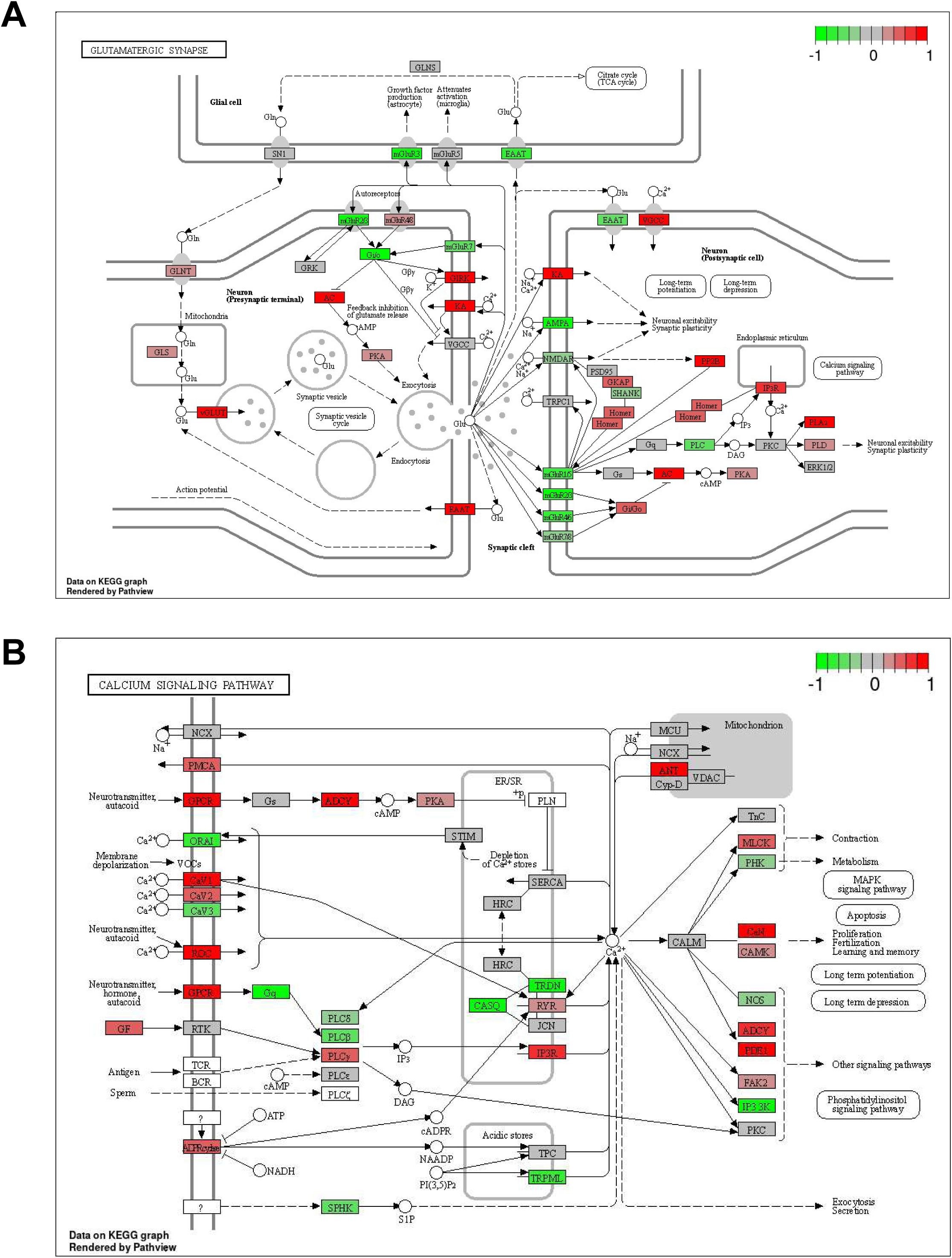
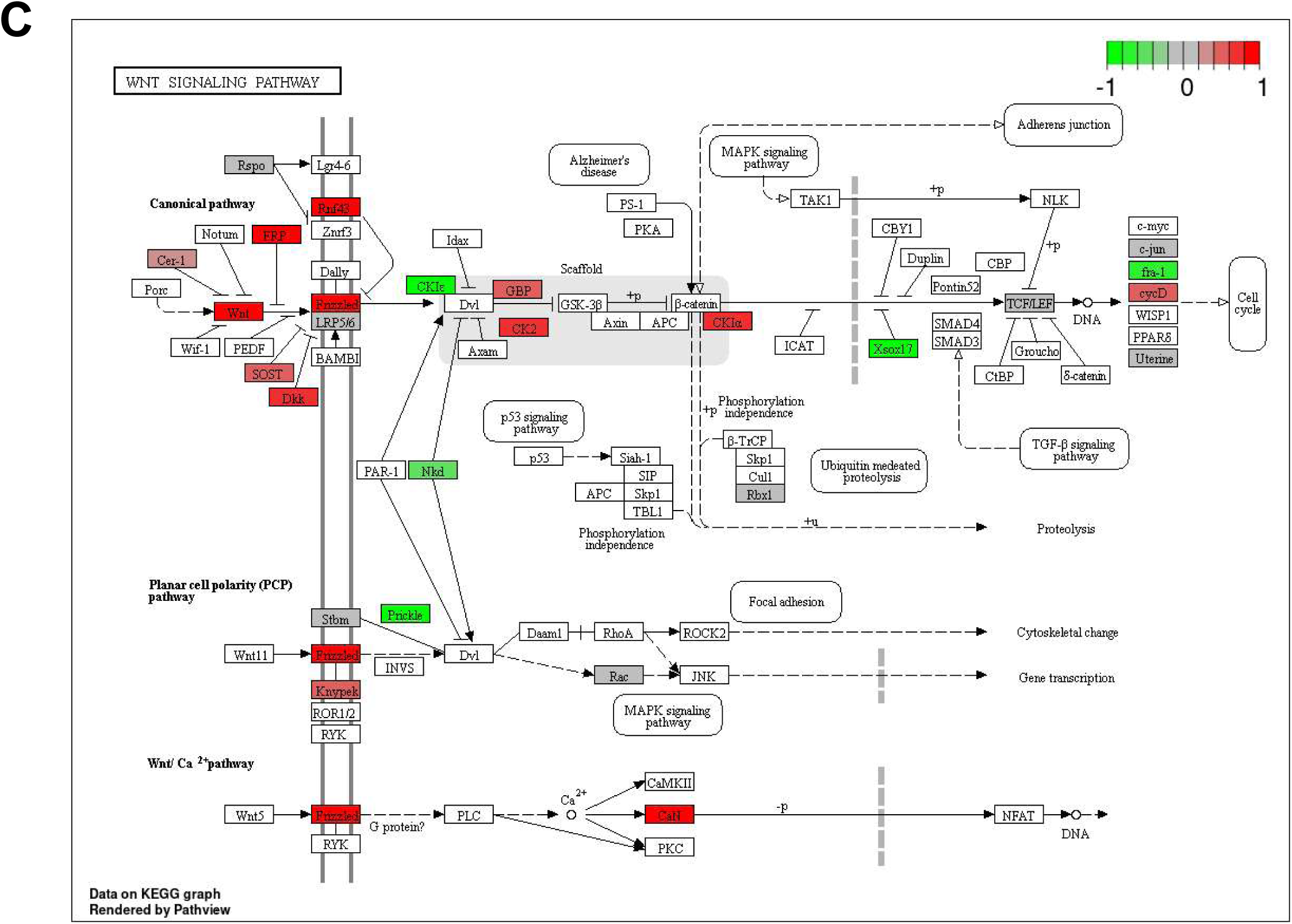
KEGG pathway enrichment analysis for glutamatergic synapse and calcium signalling pathways. Red icons indicate upregulated and green icons indicate downregulated in mutant compared to control organoids with respect to (A) glutamatergic synapse, (B) calcium signalling and (C) WNT signalling pathways.

## SUPPLEMENTAL TABLES

**Table S1.** Summary of iPSC clones isolated after reprogramming of patient PBMCs from a family with a heterozygous deletion of exon 9 of *HNF1B*.

| Line designation | Mutation | Clinical presentation | Relationship | No. of Clones isolated |
| --- | --- | --- | --- | --- |
| TF171 | HNF1B Mutation: c.1654-?_1674+?del which causes a deletion of exon 9. | Dysplastic kidneys detected antenatally | son | 6 |
| TF172 | HNF1B Mutation: c.1654-?_1674+?del which causes a deletion of exon 9. | Dysplastic kidneys detected antenatally | son | 8 |
| TF173 | No mutation | Healthy | mother | 8 |

**Table S2.**
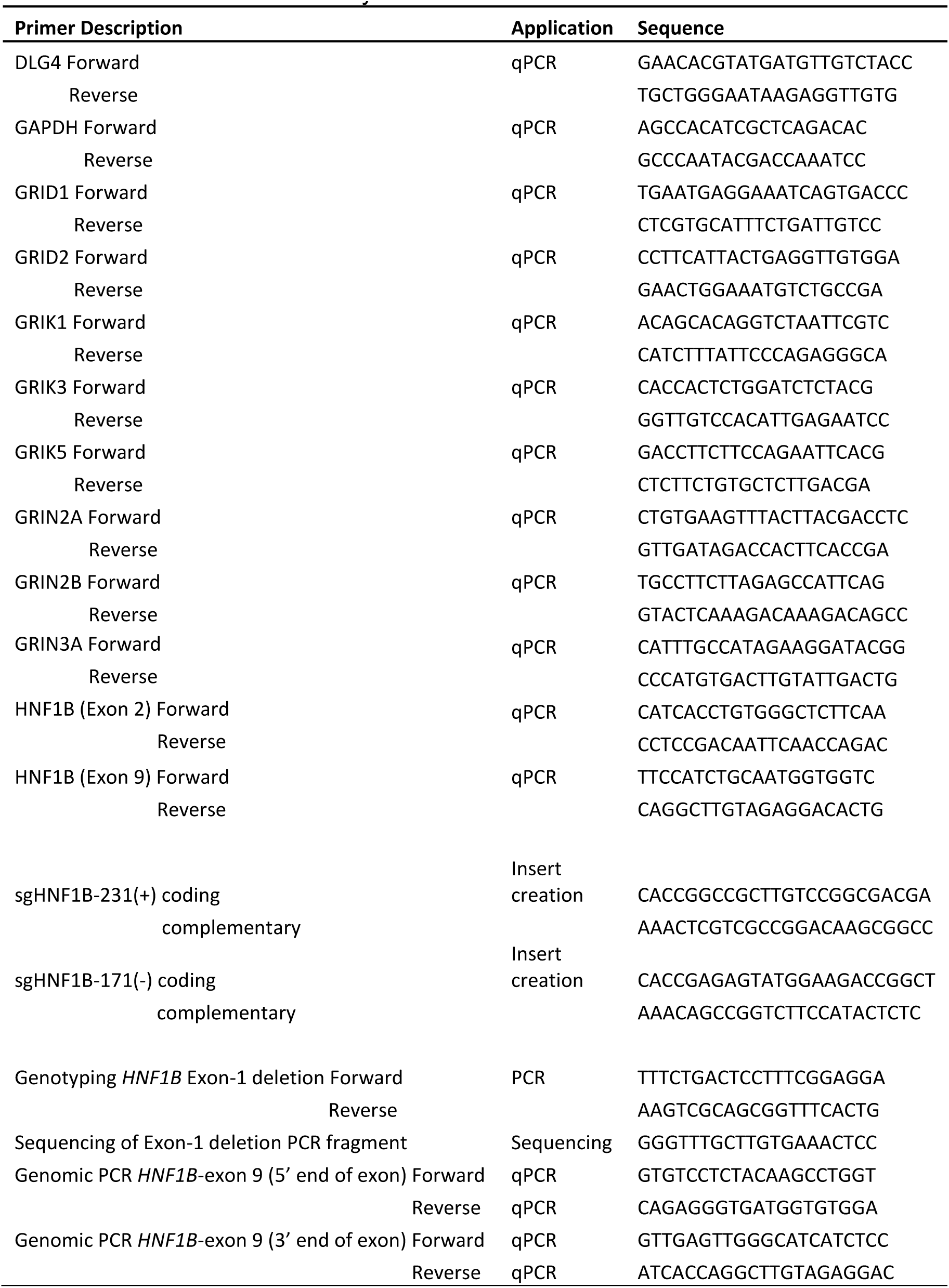
Primers used in the study

**Table S3.** Antibodies and lectins used in immunohistochemistry (IHC), immunocytochemistry (ICC) and western blot (WB) experiments.

| Primary Antibody | Host | Source | Catalogue # | Application | Dilution |
| --- | --- | --- | --- | --- | --- |
| Activated Caspase-3 | Rabbit | Abcam | ab2302 | IHC | 1:250 |
| BrdU | Rabbit | BioRad | AHP2405 | IHC | 1:100 |
| CUBN | Goat | Santa Cruz | sc-20607 | IHC | 1:100 |
| CDH1 | Mouse | Abcam | ab76055 | IHC | 1:1000 |
|  |  |  |  | ICC | 1:200 |
| GAPDH | Rabbit | Cell Signalling | 5174 | WB | 1:1000 |
| GRIK3 | Rabbit | Thermofisher | PA5-98452 | IHC | 1:800 |
|  |  |  |  | WB | 1:1000 |
| HNF1B | Rabbit | Atlas Antibodies | HPA002083 | IHC | 1:2000 |
|  |  |  |  | ICC | 1:200 |
|  |  |  |  | WB | 1:1000 |
| Megalin | Mouse | Novus Biologics | NB110-96417 | IHC | 1:200 |
| PECAM1 (CD31) | Mouse | Cell Signalling | 3528 | IHC | 1:100 |
| SYNPO | Mouse | Santa Cruz | sc-50459 | IHC | 1:200 |
| Secondary Antibody | Host | Source | Catalogue # |  | Dilution |
| Anti-Mouse IgG AlexaFluor-594 | Donkey | Thermo Fisher Scientific | A21203 | ICC | 1:300 |
| Anti-Rabbit IgG AlexaFluor-488 | Donkey | Thermo Fisher Scientific | A21206 | ICC | 1:300 |
| Biotinylated anti-Mouse IgG | Horse | Vector Laboratories | BA-2000 | IHC | 1:400 |
| Biotinylated anti-Rabbit IgG | Goat | Vector Laboratories | BA-1000 | IHC | 1:400 |
| Biotinylated anti-Goat IgG | Horse | Vector Laboratories | BA-9500 | IHC | 1:400 |
| IRDye® 800CW Anti-Rabbit | Donkey | LI-COR Biosciences | 926-32213 | WB | 1:15000 |
| Lectins |  | Source | Catalogue # | Application | Dilution |
| Biotinylated LTL |  | 2B Scientific | B-1325-2 | IHC | 1:400 |

**Table S4.** KEGG pathway enrichment analysis at different points of organoid differentiation, comparing non-mutant and mutant organoids.

|  | KEGG ID | Description | p-adjusted | Gene count<br>(fraction of<br>total in<br>pathway) |
| --- | --- | --- | --- | --- |
| Day 12 | hsa05016 | Huntington disease | 0.004474 | 6/49 |
|  | hsa00480 | Glutathione metabolism | 0.00578 | 2/10 |
|  | hsa04723 | Retrograde endocannabinoid signaling | 0.009009 | 5/29 |
|  | hsa04022 | cGMP-PKG signaling pathway | 0.011086 | 6/30 |
|  | hsa04550 | Signaling pathways regulating pluripotency of stem | 0.018149 | 11/33 |
|  | hsa04915 | Estrogen signaling pathway | 0.02521 | 5/17 |
|  | hsa04921 | Oxytocin signaling pathway | 0.030172 | 9/24 |
|  | hsa04142 | Lysosome | 0.03125 | 2/16 |
|  | hsa04724 | Glutamatergic synapse | 0.0375 | 4/18 |
|  | hsa05012 | Parkinson disease | 0.039387 | 11/44 |
|  | hsa05010 | Alzheimer disease | 0.047826 | 7/42 |
|  | hsa04714 | Thermogenesis | 0.048458 | 11/61 |
| Day 19 | hsa00562 | Inositol phosphate metabolism | 0.002604 | 1/10 |
|  | hsa04917 | Prolactin signaling pathway | 0.002732 | 4/16 |
|  | hsa04934 | Cushing syndrome | 0.003021 | 10/33 |
|  | hsa05225 | Hepatocellular carcinoma | 0.003067 | 15/35 |
|  | hsa05226 | Gastric cancer | 0.003165 | 18/43 |
|  | hsa05224 | Breast cancer | 0.003344 | 19/45 |
|  | hsa04666 | Fc gamma R-mediated phagocytosis | 0.005208 | 2/13 |
|  | hsa05217 | Basal cell carcinoma | 0.00551 | 6/19 |
|  | hsa04916 | Melanogenesis | 0.00554 | 6/21 |
|  | hsa04310 | Wnt signaling pathway | 0.006452 | 17/44 |
|  | hsa04012 | ErbB signaling pathway | 0.006515 | 1/14 |
|  | hsa04070 | Phosphatidylinositol signaling system | 0.007813 | 1/13 |
|  | hsa04740 | Olfactory transduction | 0.008086 | 39/77 |
|  | hsa04150 | mTOR signaling pathway | 0.009202 | 6/35 |
|  | hsa04020 | Calcium signaling pathway | 0.012658 | 13/43 |
|  | hsa04714 | Thermogenesis | 0.014035 | 48/62 |
|  | hsa05218 | Melanoma | 0.016854 | 10/18 |
|  | hsa05165 | Human papillomavirus infection | 0.017241 | 17/61 |
|  | hsa04530 | Tight junction | 0.019293 | 9/29 |
|  | hsa05205 | Proteoglycans in cancer | 0.022581 | 8/44 |
|  | hsa00190 | Oxidative phosphorylation | 0.023411 | 35/48 |
|  | hsa04976 | Bile secretion | 0.042071 | 5/13 |
|  | hsa03010 | Ribosome | 0.04908 | 27/38 |
| Day 25 | hsa00190 | Oxidative phosphorylation | 0.002257 | 39/48 |
|  | hsa03010 | Ribosome | 0.004484 | 34/39 |
|  | hsa04024 | cAMP signaling pathway | 0.004535 | 10/58 |
|  | hsa04714 | Thermogenesis | 0.006726 | 41/62 |
|  | hsa05010 | Alzheimer disease | 0.008696 | 33/43 |
|  | hsa04923 | Regulation of lipolysis in adipocytes | 0.01073 | 2/13 |
|  | hsa05140 | Leishmaniasis | 0.010846 | 8/17 |
|  | hsa05164 | Influenza A | 0.011062 | 13/38 |
|  | hsa04932 | Non-alcoholic fatty liver disease (NAFLD) | 0.011211 | 24/39 |
|  | hsa04917 | Prolactin signaling pathway | 0.023861 | 2/17 |
|  | hsa00230 | Purine metabolism | 0.025263 | 7/24 |
|  | hsa04940 | Type I diabetes mellitus | 0.029279 | 5/10 |
|  | hsa04080 | Neuroactive ligand-receptor interaction | 0.030879 | 46/148 |
|  | hsa04623 | Cytosolic DNA-sensing pathway | 0.035955 | 4/11 |
|  | hsa04380 | Osteoclast differentiation | 0.037815 | 3/20 |
|  | hsa04970 | Salivary secretion | 0.037815 | 10/20 |
|  | hsa04740 | Olfactory transduction | 0.039927 | 37/76 |
|  | hsa05224 | Breast cancer | 0.043182 | 17/47 |

